# Phenotypic plasticity affects species packing in seasonally fluctuating environments

**DOI:** 10.64898/2026.09.25.754459

**Authors:** David M. Anderson, Rachel B. Potter, Colin T. Kremer

## Abstract

Environmental variability influences the evolution of phenotypic plasticity and the ecological interactions among species in communities, but it is unclear how these processes interact to affect ecological systems across scales. We investigate models describing how plasticity evolves in competitive communities exposed to seasonal variation in abiotic conditions. We conducted eco-evolutionary analyses to identify communities of coexisting species with evolutionarily stable plasticity traits. These analyses revealed that the presence of interspecific competitors constrained the evolution of plasticity in some species, which instead evolved to be specialists. Plasticity often impaired species coexistence by allowing individual species to control more niche space. Communities containing plastic species were usually less species-rich but more productive and less variable. Similar results were obtained when plasticity rate was evolutionarily labile, but plasticity rate altered the patterning of temporal niche partitioning. This work shows how plasticity and competition combine to shape ecological and trait responses to environmental variability.

## Introduction

Temporal variability in the abiotic environment impacts ecological systems across levels of organization, from the phenotypes expressed by individual organisms to the composition and dynamics of communities. At the individual level, phenotypic plasticity – the capacity of an individual (or genetically identical individuals) to express different phenotypes in different environments – is a fundamental mechanism by which organisms maintain their performance in variable environments (Scheiner 1993; Schlichting & Pigliucci 1998; Stearns 1989; West-Eberhard 2003). At the community level, shifts in species interactions and community composition in response to changes in environmental conditions is an essential process that supports biodiversity and ecosystem functioning (Chesson 1994; Vellend 2016; Yachi & Loreau 1999). Despite substantial work on how phenotypic plasticity and species interactions separately moderate individual- and community-level responses to environmental variability, it is unclear how these processes interact with one another to structure ecological systems across scales (Barbour & Gibert 2021; Hendry 2016; Miner *et al*. 2005).

Environmental variability, especially when it is predictable, is expected to favor organisms that use phenotypic plasticity to adjust their traits to match environmental fluctuations (Dey *et al*. 2016; Gianoli & González-Teuber 2005; Lande 2014; Leung *et al*. 2020; Morgan *et al*. 2022; Schaum *et al*. 2018; Scheiner 1993). The scope for plastic phenotypic change depends both on the capacity for plasticity (the slope of the reaction norm describing how phenotypes vary with the environment, Gavrilets & Scheiner 1993; Lande 2014) and the rate of phenotypic change (Einum & Burton 2023; Fey *et al*. 2021; Padilla & Adolph 1996; Stomp *et al*. 2008). Theoretical research generally treats the plasticity rate as a fixed trait that constrains the evolution of plasticity capacity (Lande 2014; Padilla & Adolph 1996; Siljestam & Östman 2017), but recent works have proposed that plasticity rate itself may be selected for in variable environments (Dupont *et al*. 2024; Einum & Burton 2025). Investment in either plasticity capacity or rate likely incurs fitness costs, as remodeling phenotypes requires expending resources on traits or processes (e.g., genetic/sensory machinery or production of phenotypes) that could otherwise be spent on growth or reproduction (Auld *et al*. 2009; DeWitt *et al*. 1998; Siljestam & Östman 2017). However, research on the evolution of plasticity typically uses optimality approaches, in which the fitness value of investing in plasticity does not depend on the plasticity strategies of other species or genotypes in the environment (Gabriel 2005; Lande 2014; Padilla & Adolph 1996; Siljestam & Östman 2017). It is therefore uncertain how interactions among species in ecological communities may modify the evolution of plasticity.

Variability in the abiotic environment can influence interactions among competing species, leading to niche differentiation and supporting species coexistence. Species can coexist in environments with fluctuating abiotic conditions if they exhibit different responses to the abiotic factor (such that time periods supporting one species’ growth is less favorable to other species, Grover 1990) and employ some mechanism to buffer population losses during unfavorable time periods (the “storage effect,” Chesson 1994, 2000; Chesson & Warner 1981; but see Stump & Vasseur 2023). Fluctuations in resource supply can also support coexistence when species exhibit different non-linear responses to resource availability (“relative nonlinearity,” Chesson 1994; Grover 1990; Litchman & Klausmeier 2001). In communities in which coexistence is supported by such fluctuation-dependent mechanisms, resident species exhibit asynchronous population fluctuations that maintain high productivity and stability at the community level (Gonzalez & Loreau 2009; Ives *et al*. 1999; Polazzo *et al*. 2025; Yachi & Loreau 1999). Phenotypic plasticity has been hypothesized to reduce the scope for coexistence via abiotic niche partitioning because plasticity can expand a species’ fundamental niche and thereby reduce niche availability for competing species (Berg & Ellers 2010; Kremer & Klausmeier 2013). However, this hypothesis has not received significant attention, as investigations of plasticity’s effects on coexistence have focused primarily on plastic responses to biotic factors (e.g., competitors) that promote niche differentiation and coexistence (Hess *et al*. 2022; McGuinness *et al*. 2025; Pérez-Ramos *et al*. 2019; Puy *et al*. 2021; Turcotte & Levine 2016).

The adaptive evolution of phenotypic plasticity is linked to the composition and dynamics of communities via eco-evolutionary feedback loops (Hendry 2016). Shifts in species investment in plasticity can alter density-dependent interactions and community structure (Gómez *et al*. 2023; Miner *et al*. 2005; Turcotte & Levine 2016) and, in turn, these changes in the biotic environment can influence selection on phenotypic plasticity (Norberg 2004). Components of this eco-evolutionary feedback are well-studied; for example, large bodies of research evaluate the effects of plasticity on population dynamics and species interactions (in the absence of evolutionary adaptation, Fey *et al*. 2021; Reed *et al*. 2010; Rescan *et al*. 2020; Stomp *et al*. 2008), and the effects of species interactions on adaptation of environmental tolerance curves (in the absence of plasticity in environmental tolerance, Kremer & Klausmeier 2013, 2017; Miller & Klausmeier 2017; Snyder & Adler 2011). Yet, few studies consider the feedback between these ecological and evolutionary processes (but see, Chevin & Chauhan 2025) and, as such, it remains incompletely understood how it plays out to influence the evolution of plasticity and the diversity and structure of ecological communities.

Here, we investigate the evolution of phenotypic plasticity in competitive communities exposed to seasonal environmental variability. We develop a model describing the evolution of plasticity in environmental tolerance curves (Lande 2014) in communities structured by competition (Kremer & Klausmeier 2017). The seasonal (sinusoidal) environmental forcing is slow relative to species population dynamics, such that community composition can turn over within a seasonal cycle. Empirical systems that have these characteristics include micro-organisms or fast-growing plants and animals that form communities structured by seasonal succession (Mazzocchi & d’Alcalà 1995; Sasaki *et al*. 2025b; Sommer 1986) and exhibit plasticity in environmental tolerance curves (Anderson *et al*. 2025; Fey *et al*. 2021; Luhring & DeLong 2017; Rescan *et al*. 2022; Roeder *et al*. 2025; Sasaki *et al*. 2025a; Van Baelen *et al*. 2024). We address the following questions:

1) How do competitive interactions affect the evolution of plasticity?
2) How does the evolution of plasticity influence species coexistence, community productivity, and community temporal variability?
3) How do these effects depend on whether investment in plasticity is controlled by plasticity capacity or rate?

To answer these questions, we manipulate the presence of competitive interactions, and the presence and mode of phenotypic plasticity, and evaluate the evolution of plasticity and community properties. We find that competitive communities can support species with plasticity strategies that differ markedly from the optimal strategies observed in the absence of competition, and that plasticity impairs species coexistence but improves community productivity and stability.

## Model and Methods

### Model structure

We present a model describing the evolution of phenotypic plasticity in a competitive community inhabiting a temporally variable environment. The model of plasticity evolution builds on past work by Lande (2014). We begin by defining the per-capita population growth rate (i.e., fitness, *g_i_*) of species *i* as:

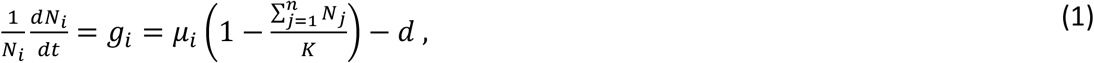

where *μ_i_* is species *i*’s intrinsic growth rate and *d* is a constant density-independent mortality rate shared by all species. The growth rate of any species *i* declines as the total density of all *n* species in the community approaches the carrying capacity *K*. The intrinsic growth rate *μ_i_* of species *i* depends on its’ strategy for shifting its’ phenotype *Z_i_*(*t*) in response to changes in abiotic environmental conditions *ε*(*t*):

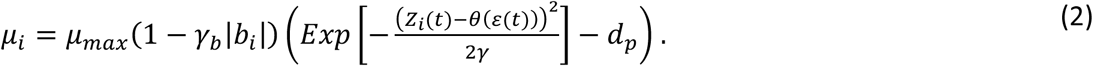

Here, *μ_i_* decreases as the optimal phenotype *θ*(*ε*(*t*)) in the environment *ε*(*t*) diverges from its current phenotype *Z_i_*(*t*) (where *γ* controls the strength of selection on these mismatches, Fig. 1A-B), and as investment in plasticity *b_i_* increases (*γ_b_* determines the strength of this cost to plasticity, Fig 1C). The optimal phenotype is linearly related to the environment following *θ*(*ε*(*t*)) = *A* + *Bε*(*t*) (Lande 2014); we simplify this expression by taking *A* = 0 and *B* = 1 and hereafter refer to *θ*(*ε*(*t*)) = *ε*(*t*) as the environment. The maximum intrinsic growth rate is *μ_max_*, and *d_p_* is an additional fitness penalty that is incurred by species investing less in plasticity when they are in extreme environments (when *Z_i_*(*t*) is very different from *θ*(*ε*(*t*)), see Fig. S1 and Supplemental Methods S1.3).

**Fig. 1.**
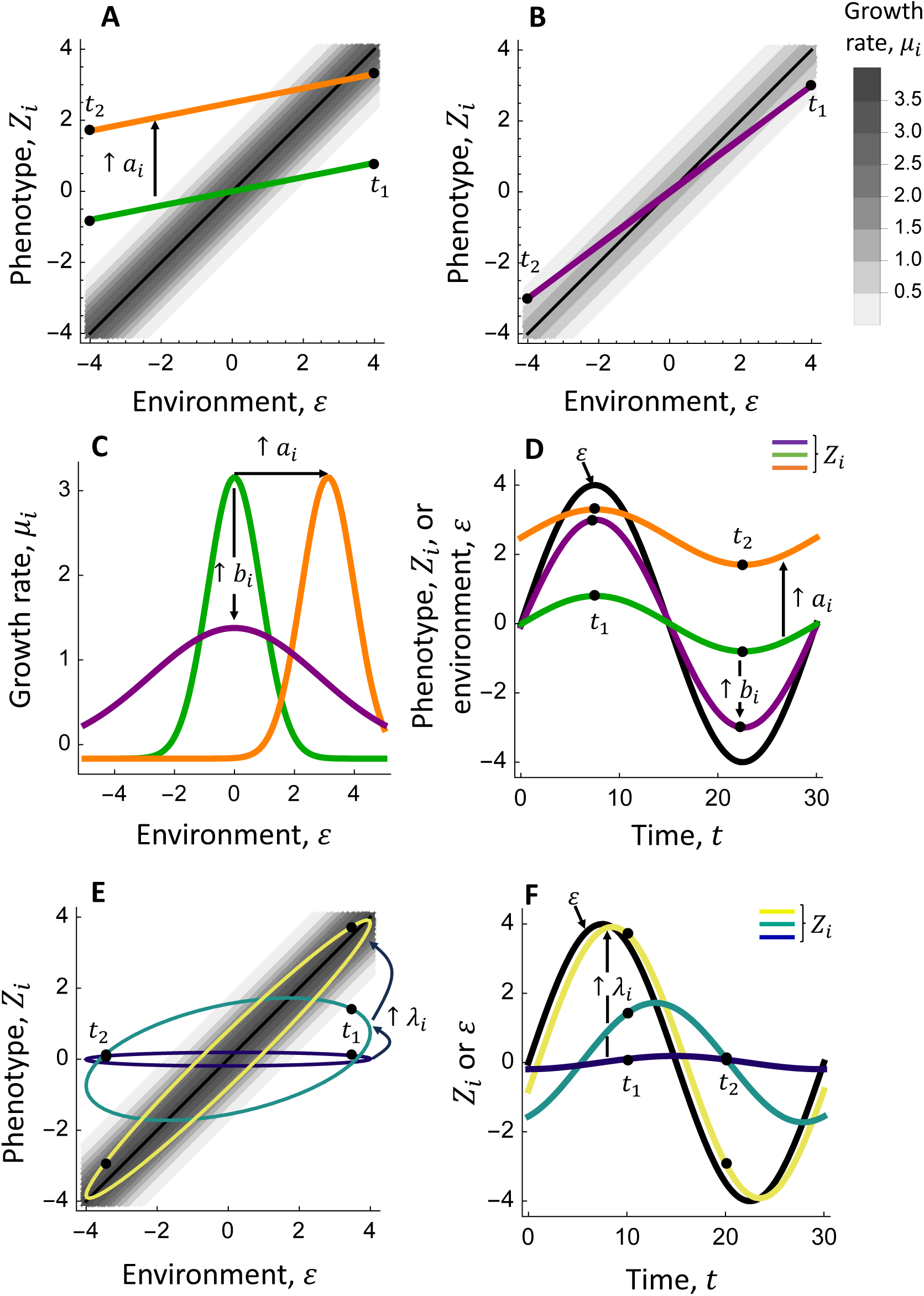
Species-specific differences in phenotypic plasticity influence growth rate and phenotype dynamics in our model (Eqn. 1-4). **A-D** Changing the intercept of the reaction norm (e.g., from *a_i_* = 0 to *a_i_* = 2.5 in **A**) shifts the environmental preferences – the environment maximizing growth rate *μ_i_* (**C**, contrast green and orange curves) – and the average phenotypes expressed across environmental cycles (**D**, compare green and orange lines). Increasing the slope of the reaction norm *φ_i_*(*ε*) = *a_i_* + *b_i_ ε* (the colored lines in **A-B**) from *b_i_* = 0.2 (**A,** green line) to *b_i_* = 0.75 (**B**, purple line) leads to a reduction in the maximum growth rate *μ_i_*, but increases the growth rate at environmental extremes (**C,** contrast green and purple curves) and leads to phenotype *Z_i_* dynamics that more closely track fluctuations in the environment *ε* (**D,** compare green and purple lines). Panels **A-D** assumes that the acclimation rate is sufficiently high (*λ_i_* = 10) that phenotype dynamics are constrained to the reaction norm. **E-F** Increasing the acclimation rate *λ_i_* – a trait that may determine investment in plasticity – from *λ_i_* = 0.01 (blue line) to *λ_i_* = 0.1 (green line) to *λ_i_* = 1 (yellow line) leads to phenotype dynamics that more closely track changes in the environment. Note that increasing acclimation rate also comes at a cost of decreased maximum growth rate (see Eqn. S1), but this is not shown here. The ellipses in panel **E** show combinations of the environment *ε* and the phenotype *Z_i_* at various points in time along the cycle in **F**, and the black points labeled *t*_1_ and *t*_2_ are included to help map sets of {*ε*, *Z_i_*} in **E** to corresponding times in **F** (these points are also included in **A-B, D**). Parameter values are *μ_max_* = 4, *γ* = 0.5, *γ_b_* = 0.85, *d_p_* = 0.05, *τ* = 30 and *λ* = 10 (in **A-D**) or *b* = 1 (in **E-F**).

Each species’ phenotype *Z_i_*(*t*) undergoes continuous, reversible development in response to changes in the environment *ε*(*t*) following (Lande 2014):

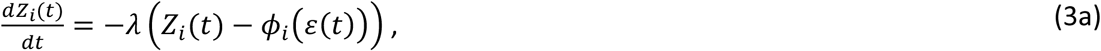

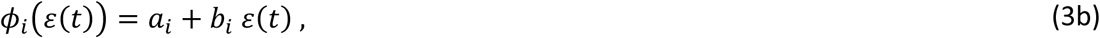

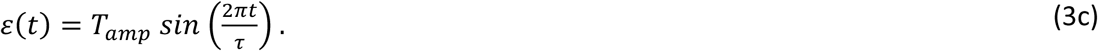

The phenotype *Z_i_*(*t*) tends towards the fixed reaction norm *φ_i_*G*ε*(*t*)H – which specifies the long-term target phenotype in the environment *ε*(*t*) – at a rate determined by the acclimation rate *λ*. The height and slope of the reaction norm of species *i* are set by *a_i_* and *b_i_*, respectively (Fig. 1A-B). The environment cycles sinusoidally with amplitude *T_amp_* and period *τ*. We solve Eqn. 3a for *Z_i_*(*t*), which describes the cycle that phenotypes settle on after long-term exposure to the environmental cycle:

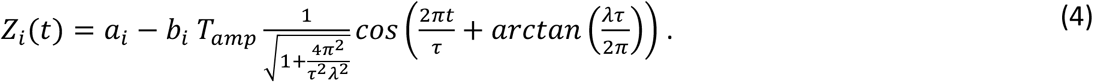

Eqn. 4 reveals that the reaction norm slope *b_i_* and the acclimation rate *λ* have different effects on phenotype dynamics: *b_i_* only influences the amplitude of the phenotype cycle (Fig. 1D), whereas *λ* additionally affects the phase of the phenotype cycle (Fig. 1F).

Species differ in two traits that determine their plasticity strategy. First, species may vary in their reaction norm intercept *a_i_*, which shifts their environmental preferences and, importantly, introduces a mechanism of niche segregation (Fig. 1A, C-D). Second, species may differ in their investment in plasticity. We consider the evolution of two plasticity mechanisms: the reaction norm slope *b_i_* or the acclimation rate *λ_i_*. Note that Eqns. 1-4 present the model in which *b_i_* evolves, whereas *λ* is fixed and thus not indexed by *i* (Supplementary Information S1.1 contains the model of *λ_i_* evolution, Eqns. S1-S3). Increases in *b_i_* (Fig. 1A-B) or *λ_i_* (Fig. 1E) come at a cost of reduced maximum growth rate (notice that growth rates are lower in Fig. 1B than 1A) but makes it easier to track the environment (Fig. 1D, F).

Eqns. 1-4 are a modified version of Lande’s (2014) model. The key change we make is to consider Gaussian rather than quadratic penalties for phenotype-environment mismatches, which reduces the negative effects of large mismatches (Fig. S1) and thereby buffers population growth during unfavorable time periods (a prerequisite for coexistence via the storage effect, Chesson 1994). Many organisms employ processes such as dormancy, torpor, or other mechanisms that reduce the negative effects of exposure to extreme environments. See Supplementary Methods S1.3 for additional details.

### Trait evolution

We evaluate trait evolution in the absence and presence of competitive interactions. In the absence of competition, the per-capita population growth rate of any species *i* is not influenced by density-dependence (i.e., 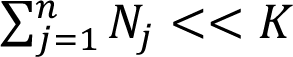 in Eqn. 1). In this case, the fitness of species *i* with traits {*a_i_*, *b_i_*} (or {*a_i_*, *λ_i_*}) is determined by the average per-capita population growth rate over the environmental cycle:

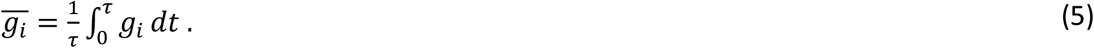

The optimal traits are those that maximize 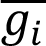 when 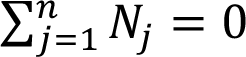. This optimality approach is commonly used to model plasticity evolution (e.g., Lande 2014), and generally does not yield diversification (Klausmeier *et al*. 2020).

To model trait evolution in the presence of competition, we use adaptive dynamics (Geritz *et al*. 1998). This approach assumes that ecological dynamics are rapid relative to evolutionary dynamics so that, if a mutant arises and replaces the resident, the system reaches its new ecological attractor before new mutants arise. We identify “Evolutionarily Stable Strategies” (ESSs, Smith & Price 1973) or “Evolutionarily Stable Communities” (ESCs, Edwards *et al*. 2018; Geritz *et al*. 1997). An ESS is a species that cannot be invaded by mutants with different traits; similarly, an ESC is a set of species that is not invasible.

To identify an ESC, we find the ecological attractor for a community with fixed traits, and then assess its evolutionary stability. At the ecological attractor, all species have zero net growth over the environmental cycle. For a community of *n* species with traits 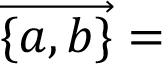 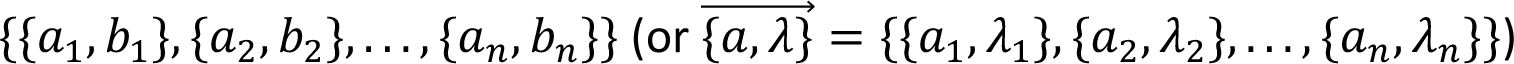 and initial (non-zero) population densities 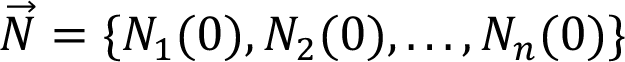 at time *t* = 0, an ecological attractor satisfies 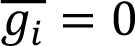 (see Eqn. 5) for each species (Kremer & Klausmeier 2017). Evolutionary stability is evaluated by assessing selection on each resident species’ traits – i.e., the slope of the relationship of fitness to trait values for an invader with traits {*a_inv_*, *b_inv_*} (or {*a_inv_*, *λ_inv_*}) near each resident (with traits 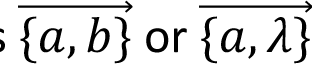):

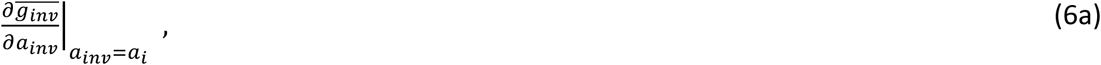

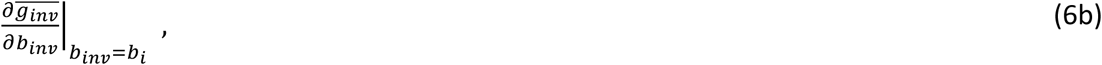

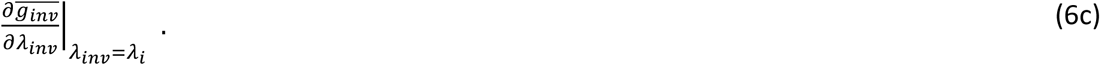

Eqns. 6 quantify selection on each trait: positive (negative) values indicate selection towards larger (smaller) trait values. When Eqns. 6 equal zero for all species in the community, the system is at evolutionary equilibrium (a singular strategy, Geritz *et al*. 1998, 2016).

A singular strategy is locally evolutionarily stable – i.e., an ESS – if mutants with trait values {*a_inv_*, *b_inv_*} (or {*a_inv_*, *λ_inv_*}) slightly larger and smaller than those of the resident(s) 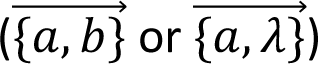 have a negative net fitness 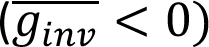 at the resident’s ecological attractor (i.e., there is stabilizing selection on the resident’s trait values). We identify globally stable ESSs – those in which all mutants, including those with traits very different from those of the resident(s), have negative invasion fitness. If a singular strategy is not locally or globally stable, the community may support an additional species. In this case, we incorporated additional species into the community and repeated eco-evolutionary analyses (Kremer & Klausmeier 2017) until we identified a community of species with trait values that are locally and globally stable (i.e., an ESC, Edwards *et al*. 2018). Hereafter, we refer to all evolutionarily stable communities, including those with only one species, as an ESC.

### Analysis

To address our questions, we examined how trait evolution responds to the amplitude of environmental fluctuations (*T_amp_*) under three scenarios that manipulate the presence of phenotypic plasticity and the mode of plasticity. We focus on *T_amp_* because it determines the amount of environmental variability, which strongly influences phenotypic plasticity (Lande 2014; Scheiner 1993) and community diversity (Kremer & Klausmeier 2017). Our three scenarios are:

1. We assessed the effect of *T_amp_* on the evolution of reaction norm traits {*a_i_*, *b_i_*} in the absence and presence of competition. The acclimation rate was set to a high value (*λ* = 10; see Fig. 1D), so that it did not limit plasticity. We considered three costs of plasticity: *γ_b_* = 0.8, 0.85, 0.9 (results for *γ_b_* = 0.8 & 0.9 are in Supplementary Results).
2. We evaluated the evolution of traits {*a_i_*, *λ_i_*} across *T_amp_* in the absence and presence of competition. We set the reaction norm slope to *b* = 1 (corresponding to a reaction norm that perfectly matches the optimal environment), so that it did not limit plasticity. Three costs of plasticity were considered: *γ_b_* = 0.3, 0.5, 0.7 (*γ_b_* = 0.3 & 0.7 are in Supplementary Results).
3. We assessed the evolution of the reaction norm intercept *a_i_* (i.e., the environmental preference) in the absence of plasticity. We did so by setting *b_i_* = 0, in which case Eqn. 4 reduces to *Z_i_* = *a_i_*.

We evaluated three properties of ESCs. First, we measured species richness as the number of coexisting species *n*. Second, we measured community productivity by calculating the average density of the community over the environmental cycle:

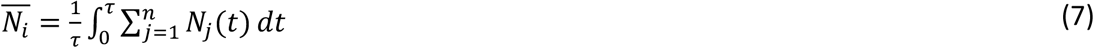

Lastly, we assessed temporal stability by calculating the variance of community density:

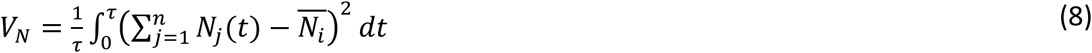

We identified ecological and evolutionary equilibria using Mathematica (Version 12.1.1.0) and the EcoEvo package. Ecological attractors and ESCs (Eqn. 1) could not be solved analytically and were therefore identified numerically using root finding methods. We used bifurcation continuation techniques to track equilibria across amplitudes (Kremer & Klausmeier 2017).

## Results

### Effects of competition on the evolution of plasticity

The evolutionary response of reaction norm traits to environmental fluctuation amplitude depends on the presence of competitive interactions. Absent competition, increases in cycle amplitude select for greater investment in phenotypic plasticity (i.e., a steeper reaction norm slope *b_i_*, Fig. 2A). This result occurs because plasticity helps species avoid the large fitness losses incurred by phenotype-environment mismatches at high amplitudes. For instance, notice that a species with the optimal *b_i_* = 0.35 at *T_amp_* = 1.5 (Fig. 2B) has relatively low fitness at *T_amp_* = 3 (Fig. 2C) because it often experiences large mismatches between its phenotype and the environment (Fig. 2D, 2G). Increased investment in plasticity (e.g., to *b_i_* = 0.72, as in the purple species, Fig. 2E) improves the match between phenotype and environment (Fig. 2G). In this case, the benefit from improved phenotype-environment matching outweighs the cost of plasticity and ultimately increases net fitness (Fig. 2C). At high amplitudes, the optimal investment in plasticity approaches the maximum investment in plasticity, *b_i_* = 1 (where the reaction norm matches the optimum).

**Fig. 2.**
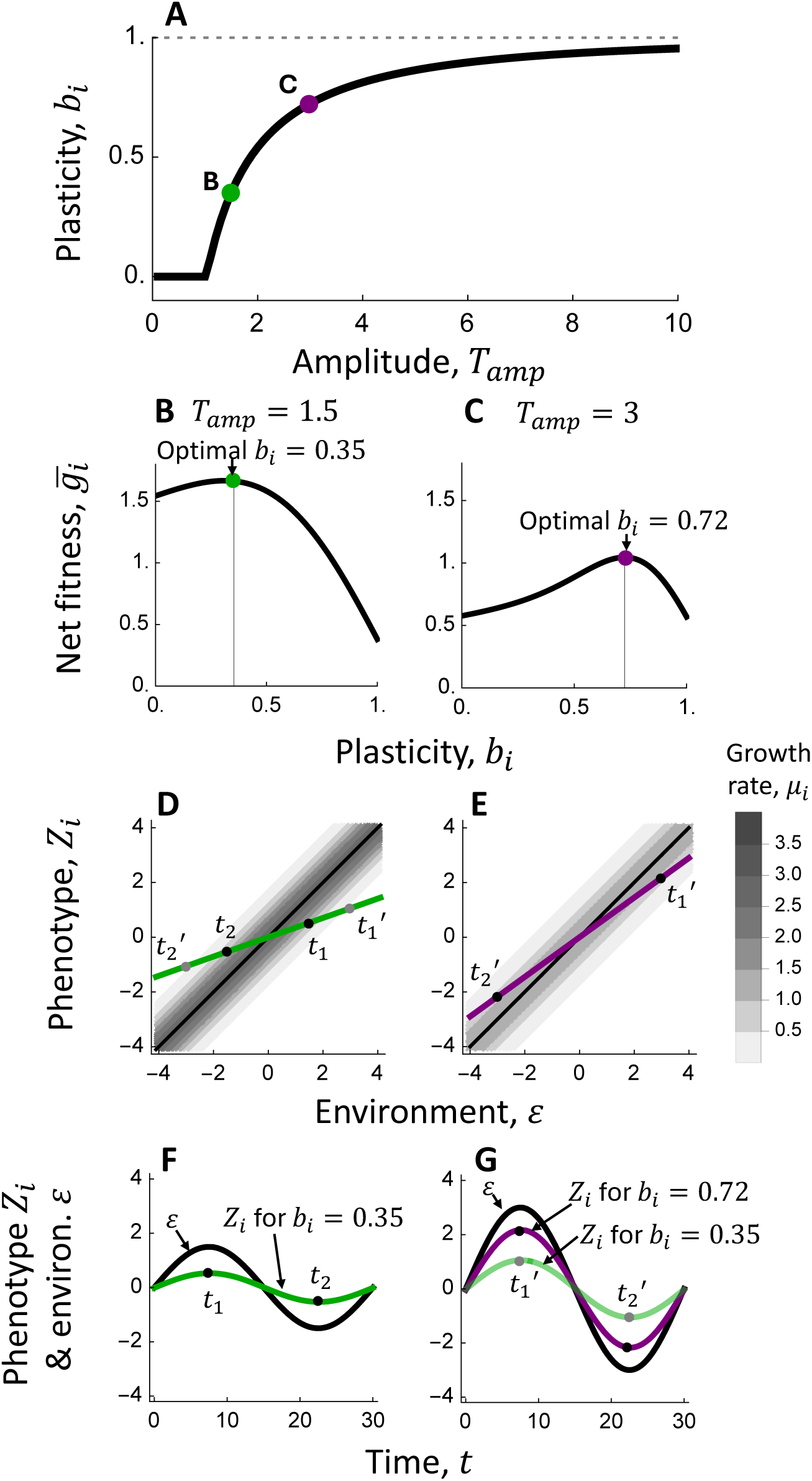
In the absence of competition, the optimal investment in phenotypic plasticity increases with the amplitude of environmental fluctuations. **A** The reaction norm slope *b_i_* maximizing the net fitness per period (when *N_i_* << *K*) increases with *T_amp_*, and at high amplitude approaches *b_i_* ≈ 1 (the dashed line, corresponding to a reaction norm that matches the optimal environment). In all cases, the optimal reaction norm intercept is *a_i_* = 0 (see Fig. S2B1). The indicated points are considered in more detail in **B-G**. **B-C** Effects of varying investment in plasticity on the net fitness per period when *T_amp_* = 1.5 and *T_amp_* = 3. **D-E** Reaction norms and growth rate *μ_i_* contour plots for the optimal traits at *T_amp_* = 1.5 and *T_amp_* = 3, which are *b_i_* = 0.35 and *b_i_* = 0.72, respectively. Note that the cost of plasticity leads the maximum growth rate *μ_i_* to be lower for the species that invests more in plasticity (shown in **E**). The points *t*_1,2_ and *t*_1,2_^E^ indicate sets of phenotypes and environments at two time points during cycles of amplitude *T_amp_* = 1.5 and *T_amp_* = 3, respectively, shown in **F-G**. **F-G** Phenotype-environment dynamics for the optimal species at *T_amp_* = 1.5 and *T_amp_* = 3. Phenotypic dynamics for the optimal species at *T_amp_* = 1.5 (transparent green line) are also displayed at *T_amp_* = 3 in panel **G**, to illustrate the relatively large phenotype-environment mismatches that arise from low investment in plasticity at this amplitude. Parameter values are *μ_max_* = 4, *γ* = 0.5, *γ_b_* = 0.85, *d_p_* = 0.05, *τ* = 30 and *λ* = 10.

In competitive communities, some species evolve very different plasticity traits than those that are optimal in the absence of competition (Fig. 3). At low amplitudes, a single plastic species maintains consistently high population densities (Fig. 3F) and outcompetes other strategies (Fig. 3C). At intermediate amplitudes, that species is no longer able to fully fill the available abiotic niche space (Fig. 3J) and maintain sufficiently high densities (Fig. 3G) to outcompete all other strategies. This leads to the emergence of two specialist species that do not invest in plasticity (orange, blue species, Fig. 3D) and instead capitalize on periods of time in which the environment is at relatively high or low values (Fig. 3J, G). These three species – the plastic generalist and two non-plastic specialists – form an evolutionarily stable community (Fig. 3D). At high amplitudes, the two non-plastic specialist species go extinct and the environment is again filled only by a single plastic species (Fig. 3A). This occurs because high amplitudes lead the plastic species to invest substantially in plasticity (and hence fill up a large portion of available niche space, Fig. 3K), and high amplitudes lead to larger environmental fluctuations that reduce the fitness value of specializing.

**Fig. 3.**
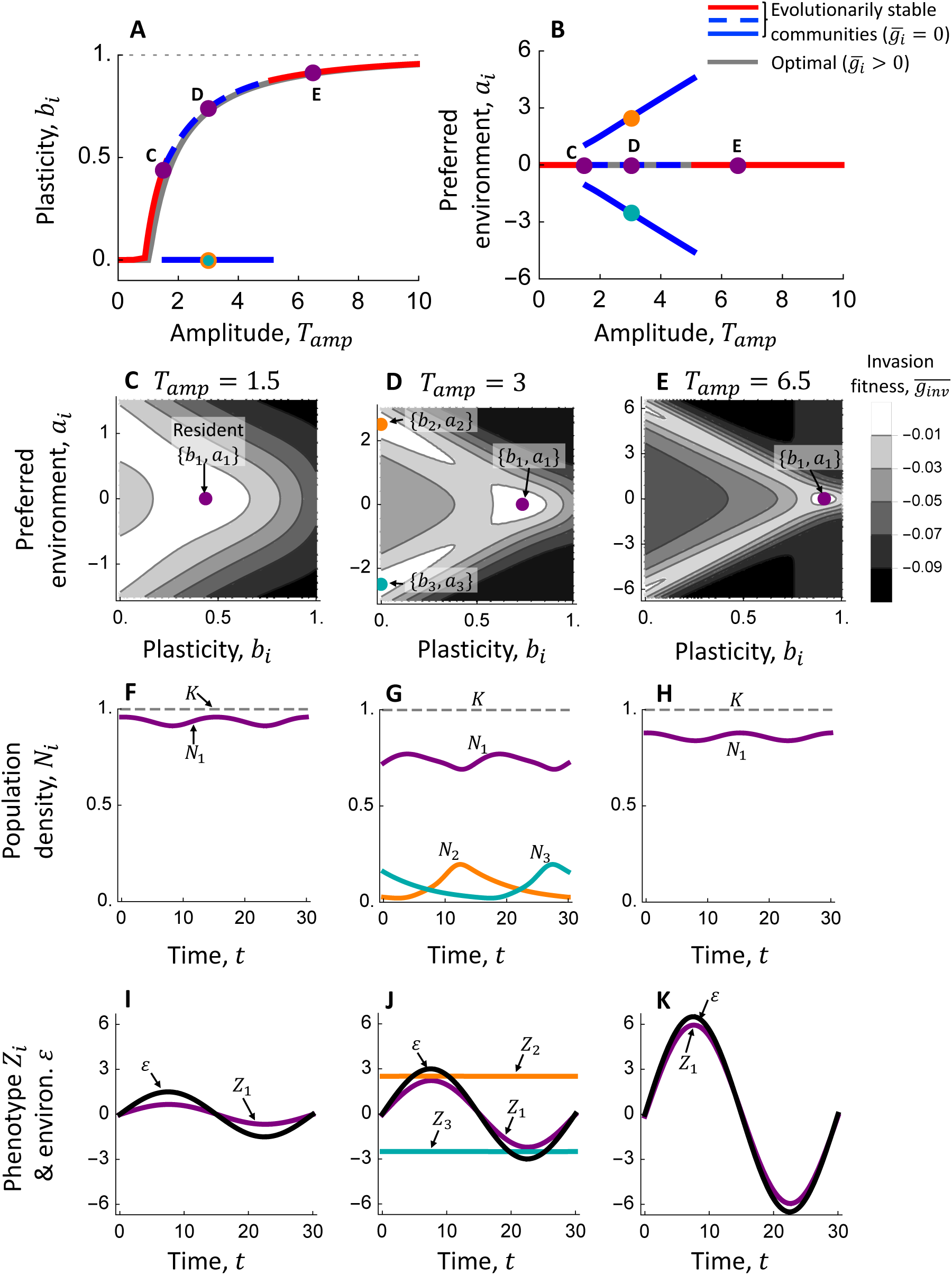
Effects of the amplitude of environmental fluctuation on the evolutionarily stable investment in plasticity *b_i_* (A) and environmental preferences *a_i_* (B). **A-B** Evolutionarily stable communities comprising species with sets of reaction norm traits 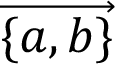 across gradients in fluctuation amplitude. The color of ESCs change with changes in the species richness of the community, and the linetypes are varied in multispecies ESCs to map the traits in **A** to those in **B**. The solid gray line shows the optimal traits found in the absence of competition (Fig. 2A), and the dashed gray line shows *b_i_* = 1 (corresponding to a reaction norm slope that matches the optimal environment). The points indicated in these panels are explored in more detail in **C-K**. **C-E** Invasion fitness for rare mutants that vary in *b_i_* and *a_i_* at the ecological attractor for the resident species or community with traits indicated by the colored points. All invaders have a negative invasion fitness, and hence the resident traits constitute ESSs or ESCs. **F-H** The ecological attractors for the ESCs at *T_amp_* = 1.5, *T_amp_* = 3, and *T_amp_* = 6.5. **I-K** Phenotype-environment dynamics for the ESCs at *T_amp_* = 1.5, *T_amp_* = 3, and *T_amp_* = 6.5. Parameter values are *μ_max_* = 4, *γ* = 0.5, *γ_b_* = 0.85, *d_p_* = 0.05, *d* = 0.1, *K* = 1, *τ* = 30 and *λ* = 10.

The cost of plasticity (*γ_b_*) influences the optimal investment in plasticity and the diversity of ESCs (Supplementary Information, S2.1). In the absence of competition, increasing costs generally select for reduced investment in plasticity (decreased *b_i_*), and ultimately species that do not invest in plasticity at all (*b_i_* = 0) but instead specialize (*a_i_* ≠ 0, Fig. S3). In the presence of competition, increasing plasticity costs generally increase the diversity of ESCs by reducing the amount of niche space that a species investing in plasticity could control (Fig. S3). Notably, at a low cost of plasticity, diversification does not occur at all and the community consists of only a single species at all amplitudes (Fig. S3A, Supplementary Information, S2.1).

### Effects of plasticity type: acclimation rate vs reaction norm slope

Competition also influences plasticity evolution when the acclimation rate *λ_i_* is evolutionarily labile and controls the scope for phenotypic plasticity (Fig. 4). Without competition, increases in environmental fluctuation amplitude often leads to increases in the optimal investment in plasticity *λ_i_*, as this helps species avoid phenotype-environment mismatches at higher amplitudes (Fig. 4A, Supplementary Information S2.2, Fig. S4). At sufficiently high amplitudes (e.g., *T_amp_* > ∼6), however, the optimal strategy involves reducing investment in plasticity and instead specializing on peaks or troughs in the environmental cycle (Fig. 4A-B, Supplementary Information S2.2, Fig. S4). These two strategies (specializing on peaks or troughs) are equally fit in our model because of the perfect symmetry of environmental tolerance curves (Fig. 1C) and the bimodal distribution of environmental states in sinusoidal cycles (Kremer & Klausmeier 2017).

**Fig. 4.**
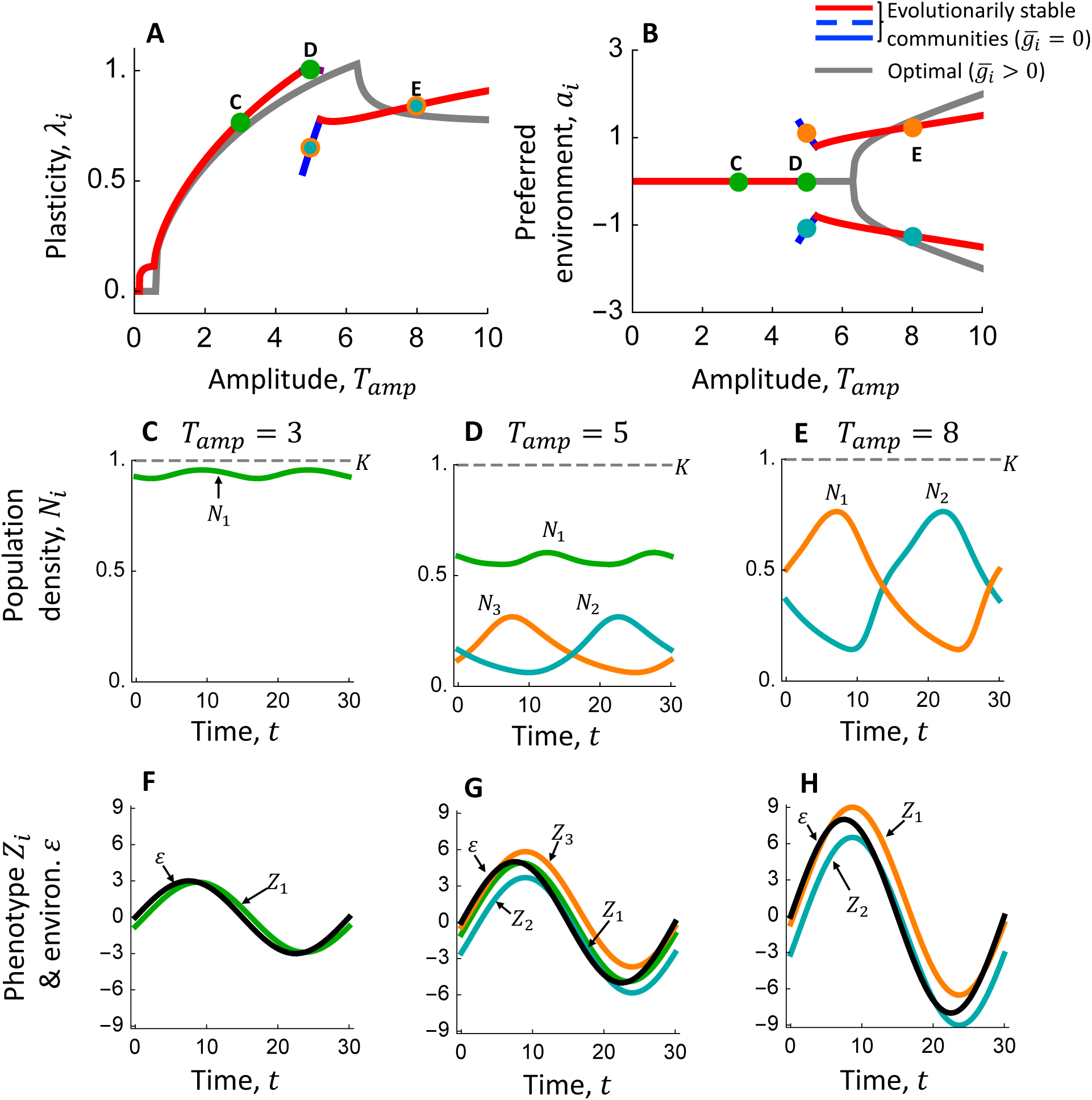
Evolutionarily stable investment in plasticity *λ_i_* (A) and environmental preferences *a_i_* (B) across a gradient in fluctuation amplitude. **A-B** Effect of *T_amp_* on evolutionarily stable communities consisting of species that differ in environmental preferences and acclimation rate 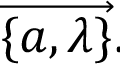. The color of ESCs change with changes in the species richness of the community, and the gray line shows the optimal traits found in the absence of competition (more details of optimal traits are given in Supplementary Information S2.2 and Fig. S4). The points indicated in these panels are considered further in **C-H** (note that the green point at *T_amp_* = 5 covers this species’ curve). **C-E** The population dynamics (or ecological attractors) for the ESCs at *T_amp_* = 3, *T_amp_* = 5, and *T_amp_* = 8. **F-H** Phenotype-environment dynamics for the ESCs at *T_amp_* = 3, *T_amp_* = 5, and *T_amp_* = 8. Parameter values are *μ_max_* = 4, *γ* = 0.5, *γ_b_* = 0.5, *d_p_* = 0.05, *d* = 0.1, *K* = 1, *τ* = 30 and *b* = 1.

Evolution in the presence of competition leads to the emergence of ESCs containing species with traits that often differ from the optimal observed without competition. At low fluctuation amplitudes (e.g., *T_amp_* = 3), the community consists of a single plastic species (Fig. 4A-C, F). At sufficiently high amplitudes (e.g., *T_amp_* = 5), two specialist species can invade and coexist with the plastic species (Fig. 4D, G). Further increases in amplitude leads to the extinction of the plastic species (i.e., the green species in Fig. 4D) and an ESC comprising two specialist species (Fig. 4E, H). These patterns are robust to variation in plasticity costs: increasing the cost of plasticity reduces the evolutionarily stable investment in plasticity *λ_i_* and shifts the bifurcation events (i.e., species gain/loss) described above to lower amplitudes (Supplementary Information S2.3, Fig. S5).

Interestingly, the patterning of temporal niche partitioning among members of ESCs differs qualitatively between the two modes of plasticity (acclimation rate *λ_i_* in Fig. 4 vs. reaction norm slope *b_i_* in Fig. 3). Consider the 3-spp. ESC observed when the acclimation rate *λ_i_* evolves (Fig. 4D, G). Notice that the two specialists grow during intermediate environments (i.e., when *ε* ≈ 0 at times ∼0 and ∼15, see Fig. 4G). This occurs because the specialist invests only moderately in plasticity, which leads their phenotypes to lag behind the environment and ultimately achieves phenotype-environment matches during intermediate environments when *ε* ≈ 0 (Fig. 4G). This manner of temporal niche partitioning contrasts with that observed when the reaction norm slope is evolutionarily labile, in which the specialists grow during peaks and troughs (corresponding to their respective environmental preferences *a_i_*, Fig. 3G, J).

### Effects of phenotypic plasticity on community properties

Communities containing species that may be plastic (Fig. 3-4) are generally less diverse but maintain higher densities and reduced temporal fluctuations than communities containing species that do not invest in plasticity (Fig. 5-6). In Fig. 5, we present ESCs formed by species that do not invest in plasticity (*b_i_* = 0). Absent plasticity, communities reach a higher diversity – 4 species (Fig. 5A-B, Fig. 6A) – because individual species with fixed phenotypes cannot capture as much niche space as species that invest in plasticity (Fig. 5D). Strategies involving intermediate environmental preferences (i.e., with *a_i_* = 0) are also less viable – notice that no species with *a_i_* = 0 can persist at *T_amp_* > ∼3 in Fig. 5A, whereas species with *a_i_* = 0 persist at all amplitudes when *b_i_* is not fixed at 0 (Fig. 3B) and up to *T_amp_* ≈ 5 under labile *λ_i_* (Fig. 4B). Intermediate strategies are less viable because sinusoidal environments spend relatively less time at intermediate values as *T_amp_* increases (Kremer & Klausmeier 2017). At high amplitudes, communities contain 2 species that specialize in peaks and troughs of the cycle (Fig. 5D, F).

**Fig. 5.**
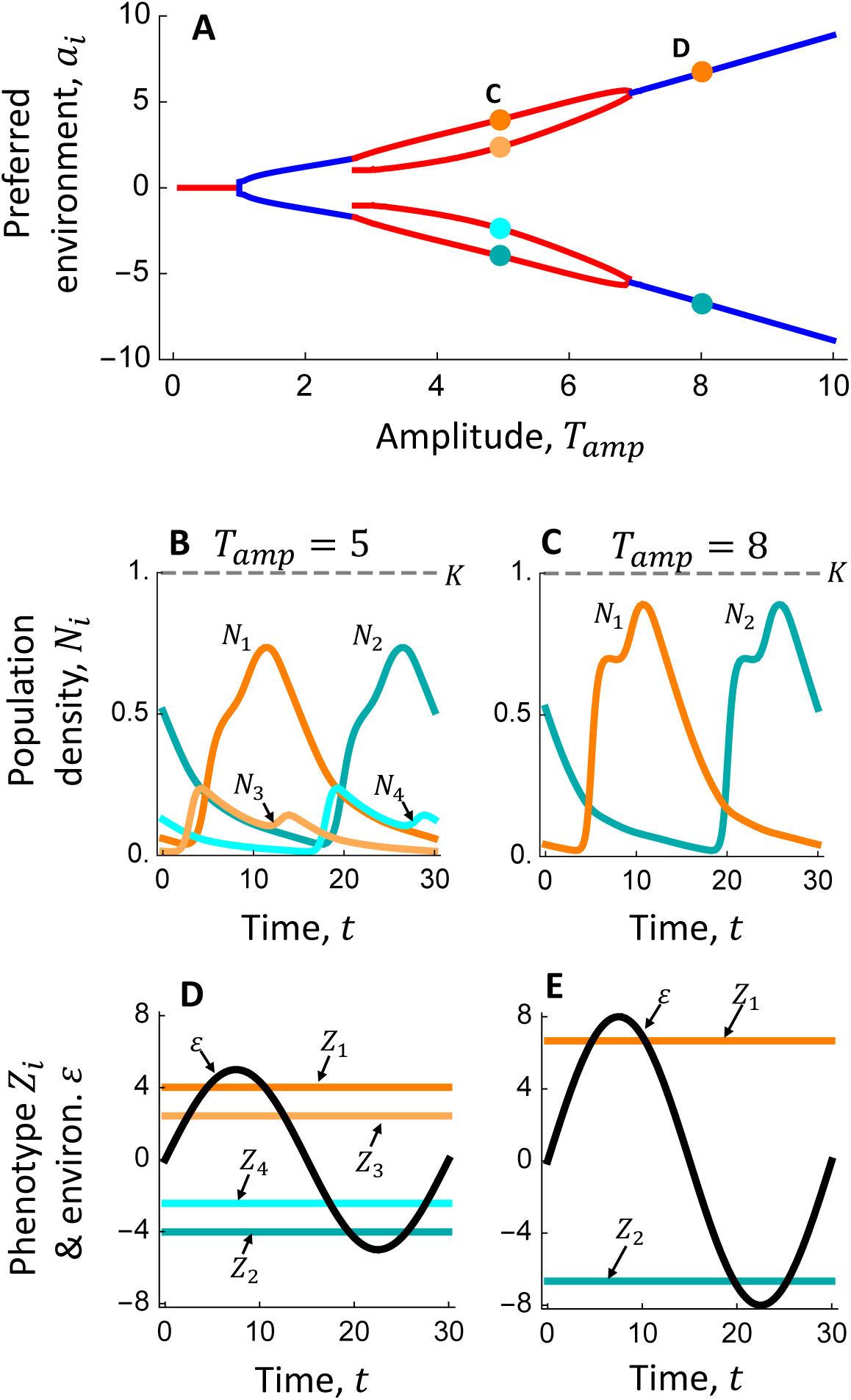
Effects of environmental fluctuation amplitude on evolutionarily stable communities in the absence of phenotypic plasticity (i.e., when *b* = 0). **A** Non-plastic species diversify only in their environmental preferences *a_i_*. The color of ESCs change with changes in species richness, and the indicated points are explored further in **B-E**. **B-C** Population dynamics and corresponding **D-E** phenotype-environment dynamics for the ESCs at *T_amp_* = 5, and *T_amp_* = 8. Because species cannot invest in plasticity in this scenario, phenotypes are constant. Parameter values are *μ_max_* = 4, *γ* = 0.5, *d_p_* = 0.05, *d* = 0.1, *K* = 1, *τ* = 30 and *b* = 0 (note that *γ_b_* vanishes in Eqn. 2 when setting *b* = 0).

**Fig. 6.**
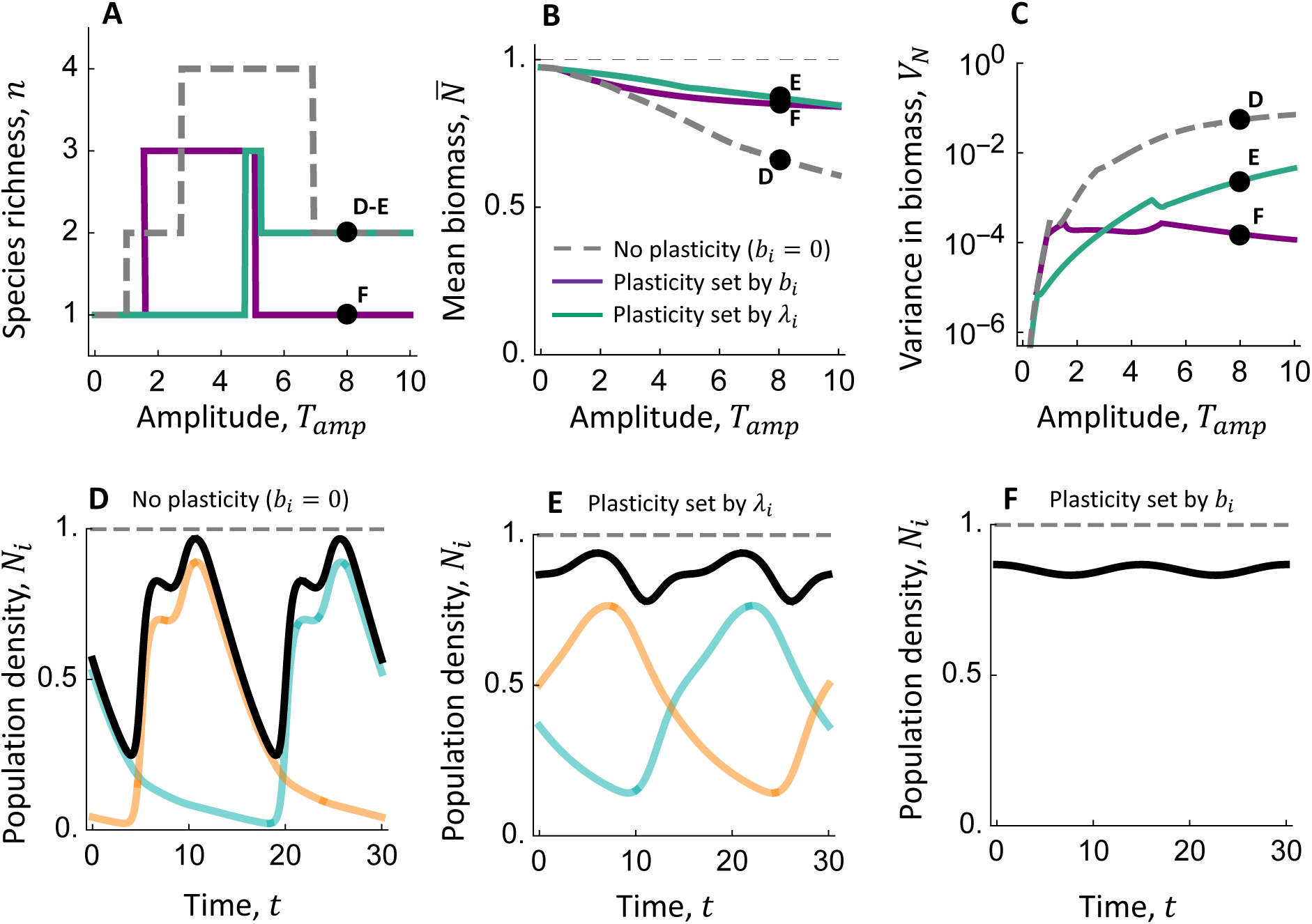
Effects of phenotypic plasticity on the response of community properties – diversity (A), mean biomass (B, Eqn. 7), and variance in biomass (C, Eqn. 8) – to the amplitude of environmental fluctuations. **A-C** These panels show effects of *T_amp_* on properties of ESCs obtained in the absence of plasticity (gray, dashed line, Fig. 5A), when plasticity is set by *b_i_* (purple line, Fig. 3A-B), and when plasticity is set by *λ_i_* (blue line, Fig. 4A-B). The black points indicate communities that will be examined in more detail in **D-F. D-F** Population and community dynamics at *T_amp_* = 8 for the three scenarios. The black line shows the total community density, the semi-transparent colored lines show individual species (note that there is only one species in **F**), and the gray dashed line shows the carrying capacity. Parameter values for each scenario can be found in the respective figure caption.

At high amplitudes, communities containing species investing in phenotypic plasticity maintain higher biomass densities (on average over the cycle, Fig. 6B) and exhibit reduced temporal fluctuations in density (Fig. 6C). This is evident by contrasting the community dynamics displayed in Fig. 6D-F. Notice that the community formed by non-plastic species exhibits large fluctuations that ultimately reduce the mean density because the community does not fully exploit niche space: it is missing a species that can grow during times in which the environment is at intermediate values (Fig. 5E). Phenotypic plasticity allows species to more efficiently fill niche space, and as such the communities in which species invest in plasticity can achieve higher densities and reduced fluctuations (Fig. 6E-F). These differences in productivity and temporal variability hold even in environments in which the non-plastic community is more species-rich (e.g., *T_amp_* = 6).

Increasing plasticity costs generally decreased the total community density across amplitudes and led to increases in the temporal variability of communities, both when the reaction norm slope and acclimation rate controls investment in plasticity (Fig. S6 & S7, respectively). This occurred because increased plasticity costs led species to reduce investment in plasticity (Supplementary Information, S2.2-S2.3), such that the community is less efficient at filling niche space (i.e., there are more time periods in which phenotype-environment mismatches reduce growth) and thus exhibits reduced productivity and increased fluctuations.

## Discussion

Our study yields three key insights into how the interplay between the adaptive evolution of plasticity and the ecological dynamics of competitive communities structures ecological and trait responses to environmental variability. First, we found that eco-evolutionary processes could select for species with plasticity traits that differed substantially from those that were optimal in the absence of competition – often including species that do not invest in plasticity at all (Fig. 3). Second, the type of plasticity (i.e., whether investment in plasticity is achieved via plasticity capacity *b_i_* or acclimation rate *λ_i_*) influenced the timing of niche partitioning among community members, which altered the patterning of diversity across fluctuation amplitudes (Fig. 4, 6). Lastly, phenotypic plasticity generally reduced the diversity of communities while also increasing their total densities and decreasing their temporal variance, as evidenced by comparisons to communities observed in the absence of plasticity (Fig. 6).

The finding that plasticity often reduced the community diversity contrasts with recent work reporting that plasticity can enhance species coexistence (Chevin & Chauhan 2025; Hess *et al*. 2022; McGuinness *et al*. 2025; Muthukrishnan *et al*. 2020; Pérez-Ramos *et al*. 2019; Puy *et al*. 2021) (but see Gibbs *et al*. 2025; Kalirad & Sommer 2024). This contrast arises from differences in how trait plasticity influences niche differentiation and species interactions. Plastic adjustment of resource-use traits induced by the presence of interspecific competitors has been found to promote coexistence by improving competitive ability (Puy *et al*. 2021) and causing niche differentiation (Pérez-Ramos *et al*. 2019). Conversely, we found that plastic responses to temporally fluctuating abiotic factors impairs species coexistence by allowing plastic species to control more of the abiotic niche, consistent with the conceptual hypothesis of Berg & Ellers (2010) (see also Kremer & Klausmeier 2013). A well-studied empirical example of this phenomenon comes from work on chromatic adaptation to light color in phytoplankton, in which a plastic species that shifts its photosynthetic pigments to match light conditions was found to competitively exclude two non-plastic species with fixed pigments in an environment with fluctuating light colors (Stomp *et al*. 2008). Taken together, this work shows that plasticity can promote or restrict species coexistence depending on how it moderates species interactions.

Our analysis highlights the importance of the acclimation rate (or plasticity rate) as a potentially evolutionarily labile trait that controls investment in plasticity. In some respects, the acclimation rate and plasticity capacity (i.e., the reaction norm slope) have similar effects on phenotypic plasticity: increases in both traits allow species to better match phenotypes to environmental fluctuations (Fig. 1, Burton *et al*. 2022; Dupont *et al*. 2024; Padilla & Adolph 1996) and, as such, we found that increases in fluctuation amplitude usually led to increased investment in both (Fig. 3-4). This result supports the hypothesis that plasticity rate evolves in more variable environments (Dupont *et al*. 2024; Einum & Burton 2023, 2025). We limited our analysis to cases in which plasticity capacity or rate were evolutionarily labile, but these traits likely coevolve and may trade-off with one another (Burton *et al*. 2020; Einum & Burton 2025). Interestingly, we found that variability in species’ plasticity rates can create more flexibility in species’ niches than variability in plasticity capacity (contrast the timing of phenotype-environment matches in Fig. 1D, F). This indicates that variability in these two traits could allow species with similar environmental preferences to access different niche spaces, thereby supporting diversification and coexistence.

We found that some species within communities did not evolve increased investment in plasticity in more variable environments (Fig. 3A), which contrasts with previous theoretical findings (Lande 2014; Scheiner 1993; Tufto 2000). There are two reasons for this discrepancy. First, our model incorporates relatively weak selection on plasticity and thereby increases the viability of non-plastic species (see Supplementary Materials S1.3). Second, we not only identified the optimal plasticity traits of a single species, but also the evolutionarily stable plasticity traits of communities of coexisting species. This latter analysis revealed species with plasticity traits that were not optimal in isolation, but supported persistence (e.g., Fig. 3D). This result could be helpful for interpreting results of meta-analyses that yield mixed support for theoretical predictions of plasticity evolution. Some studies have reported no or inconsistent effects of thermal variability in a species’ local environment on thermal plasticity capacity (e.g., Barley *et al*. 2021; Gunderson & Stillman 2015; Thomas *et al*. 2016). These negative results are typically attributed to difficulties with accounting for behavioral thermoregulation that decouples environmental observations from those that organisms experience (Gunderson & Stillman 2015) and/or physiological trade-offs that limit plasticity (e.g., between acclimation capacity and niche breadth, Barley et al. 2021). Our results suggest an additional possible mechanism: non-plastic species could be sacrificing plasticity to fill an open ecological niche.

Our study was necessarily limited in scopes, including with respect to the possible mechanisms of plasticity and timescales of evolutionary dynamics. First, in our model phenotypes can only respond to the prevailing environment, but many species use cues to shift traits in anticipation of future environmental conditions (Bernhardt *et al*. 2020; Usinowicz & O’Connor 2023). In some sense, phenotype dynamics under anticipatory plasticity may resemble those observed under fast plasticity rates, as both mechanisms help prevent phenotypes from lagging behind environmental fluctuations. However, imperfect anticipatory plasticity may create qualitatively different phenotype-environment mismatches (e.g., phenotypes over-shooting the reaction norm), which may affect eco-evolutionary outcomes. Second, we studied a form of plasticity in which trait change is reversible and may operate across generations, but many species exhibit irreversible trait plasticity that reflects the environment of development (West-Eberhard 2003). The form of plasticity may not qualitatively alter our results because it primarily influences the sensitivity of plasticity evolution to environmental predictability (Lande 2014); we studied a deterministic sinusoidal environment and hence did not consider effects of predictability. Third, we assumed that evolutionary dynamics are much slower than ecological dynamics (following adaptive dynamics theory, Geritz *et al*. 1998), yet evolution can occur on ecological timescales (Hairston Jr *et al*. 2005). Rapid evolution provides an additional mechanism by which species may track environmental fluctuations, and this mechanism can constrain the evolution of plasticity (Botero *et al*. 2015) and impair coexistence (Kremer & Klausmeier 2013).

Temporal variability in abiotic conditions structures ecological systems by shaping the expression of phenotypic plasticity by individual organisms and the competitive interactions among species in ecological communities. We investigated the interplay between these mechanisms by incorporating phenotypic plasticity into eco-evolutionary models of seasonally fluctuating environments. The results showed that communities may often harbor species with plasticity traits that are overlooked by optimality models that neglect community-level interactions, and that plasticity impairs fluctuation-dependent coexistence mechanisms. Overall, this study shows that eco-evolutionary perspectives can offer new insights into both the adaptive evolution of phenotypic plasticity, and the role of plasticity in shaping the structure and dynamics of communities.

## Supporting information

All files.

