## Supplementary material for "Phenotypic plasticity affects species packing in seasonally fluctuating environments": All files.

**Table of contents:**


- **S1:** Supplementary Methods
  - **S1.1:** Equations for model describing evolution of acclimation rate
  - **S1.2:** Table S1, describing all symbols used in the model.
  - **S1.3:** Contrasting our model with alternative frameworks. Supplementary text and Fig. S1
- **S2:** Supplementary Results
  - **S2.1:** Effects of cost of plasticity on evolution of reaction norm slope. Supplementary Text and Fig. S2-S3.
  - **S2.2:** The optimal acclimation rate in the absence of competition. Supplementary Text and Fig. S4.
  - **S2.3:** Effects of cost of plasticity on evolution of acclimation rate. Supplementary Text and Fig. S5.
  - **S2.4:** Effects of cost of plasticity on community richness, density, and stability. Supplementary Text and Fig. S6-S7.

### S1 Supplementary Methods

#### *S1.1 Acclimation rate model equations*

We present the full model equations in which the acclimation rate $\lambda_i$ determines a species' investment in plasticity. Population dynamics follow Eqn. 1 in the main text. However, the intrinsic growth rate $\mu_i$ of species $i$ is now described by:

$$\mu_i = \mu_{max}(1 - \gamma_b|\lambda_i|)\left(Exp\left[-\frac{(Z_i - \theta(\varepsilon))^2}{2\gamma}\right] - d_p\right). \quad \text{(S1)}$$

Notice here that the intrinsic growth rate $\mu_i$ now declines with increasing investment in plasticity $\lambda_i$. All other terms are as described in Eqn. 2 of the main text. Phenotype dynamics are given by:

$$\frac{dZ_i(t)}{dt} = -\lambda_i\big(Z_i(t) - \phi_i(\varepsilon(t))\big)\,, \quad \text{(S2a)}$$

$$\phi_i(\varepsilon(t)) = a_i + b\varepsilon(t)\,. \quad \text{(S2b)}$$

The solution to Eqn. S2a is:

$$Z_i(t) = a_i - b\,T_{amp}\frac{1}{\sqrt{1+\frac{4\pi^2}{\tau^2{\lambda_i}^2}}}cos\left(\frac{2\pi t}{\tau} + arctan\left(\frac{\lambda_i\tau}{2\pi}\right)\right). \quad \text{(S3)}$$

Eqn. S3 (and Eqn. 4 in the main text) was obtained by solving Eqn. S2a with initial conditions $Z_i(0) = Z_{i,0}$ and taking the limit $\lim_{t\to\infty} Z_i(t)$, which leads all terms containing $Z_{i,0}$ to vanish. In Eqns. S2-3, the acclimation rate $\lambda_i$ is now a species-specific trait and therefore indexed by $i$, whereas the reaction norm slope $b$ is fixed at a common value for all species.

*S1.2 Table describing all symbols used in the model.*

**Table S1 All symbols used in the model, along with a description and a value or range.**

| Symbol | Description | Value or Range |
|---|---|---|
| $N_i$ | Population density of species $i$ | Variable |
| $n$ | Number of species in the community | Variable |
| $g_i$ | Instantaneous per-capita population growth rate (or, equivalently, the fitness) of species $i$ | Variable |
| $Z_i(t)$ | Phenotype of species $i$ at time $t$ | Variable |
| $\varepsilon(t)$ | Environment at time $t$ | Variable |
| $a_i$ | Intercept of reaction norm of species $i$. Also referred to as the preferred environment. | Variable |
| $b_i$ or $b$ | Slope of reaction norm of species $i$ (or a fixed parameter when the acclimation rate is labile) | Variable or 1 |
| $\lambda_i$ or $\lambda$ | Acclimation rate of species $i$ (or a fixed parameter when the reaction norm slope is labile) | Variable or 10 |
| $\mu_{max}$ | Maximum intrinsic growth rate | 4 |
| $\gamma_b$ | Cost of plasticity | Varies (0.8, 0.85, 0.9 or 0.3, 0.5, 0.7) |
| $\gamma$ | Niche width | 0.5 |
| *A* | Intercept of relationship of environment to optimal phenotype | 0 |
| *B* | Slope of relationship of environment to optimal phenotype | 1 |

| | | |
|---|---|---|
| $d_p$ | Penalty for phenotype-environment mismatches | 0.05 |
| $d$ | Mortality rate | 0.1 |
| $K$ | Carrying capacity | 1 |
| $T_{amp}$ | Amplitude of environmental cycle | Varies (0-10) |
| $\tau$ | Period of environmental cycle | 30 |

*S1.3 Contrasting our model to alternative frameworks*

Our model of phenotypic plasticity builds on the model of Lande (2014). In Lande (2014), the per-capita rate of population growth in the absence of any density dependence, denoted $m$, depends on the difference between the phenotype $z_t$ and the optimal phenotype $\theta_t$ following:

$$m = m_{max} - \frac{\gamma}{2}(z_t - \theta_t)^2 - \frac{\gamma_b}{2} b^2 \,, \qquad \text{(S4)}$$

where $m_{max}$ is the maximum growth rate, $\gamma$ is the strength of stabilizing selection, $\gamma_b$ is the cost of plasticity, and $b$ is the reaction norm slope. As with our model, the phenotype varies with the environment following $z_t = a + b\varepsilon_t$, whereas the optimal phenotype in any environment is described by $\theta_t = A + B\varepsilon_t$. We here take $A = 0$ and $B = 1$ so that $\theta_t = \varepsilon_t$.

Importantly, under Eqn. S4, per-capita population growth rates are not bounded in extreme environments (when $z_t$ differs substantially from $\varepsilon_t$) – $m$ tends towards negative infinity as $\varepsilon_t$ diverges from $z_t$ (so long as $b \neq 1$) (Fig. S1A). This behavior makes it challenging for species that specialize on particular environments to persist because, during periods of time in which the species are mal-adapted (e.g., seasons in which $\varepsilon_t$ is far from $z_t$), population densities can decrease very quickly (i.e., per-capita population growth rates can be very large negative values). Many organisms are known to use strategies to buffer population growth rates in extreme environments (e.g., dormancy, torpor), which would serve to bound environmental tolerance curves at some finite value as $\varepsilon_t$ diverges from $z_t$. This behavior can be modeled with a gaussian environmental tolerance curve. For example, the right-hand side of Eqn. S4 could be exponentiated to give:

$$m = M_{max} Exp[-\frac{\gamma_b}{2} b^2] Exp[-\frac{\gamma}{2}(z_t - \theta_t)^2] \,, \quad \text{(S5)}$$

where $M_{max} = Exp[m_{max}]$. In Eqn. S5, the per-capita population growth rate now tends towards 0 as $\varepsilon_t$ diverges from $z_t$ (see Fig. S1B).

Our model (Eqn. 2 of the main text) modifies Eqn. S5 in two additional respects. First, rather than assuming that the maximum growth rate declines non-linearly with increasing investment in plasticity, as given by the term $Exp[-\frac{\gamma_b}{2} b^2]$ in Eqn. S5, we take the simpler assumption that this relationship is linear (described by $(1 - \gamma_b|b_i|)$, see Eqn. 2). Second, we incorporate an additional loss term, $d_p$, which leads growth rates to settle on lower values in extreme environments when species invest less in plasticity (see Fig. S1C). This second assumption leads the tail behavior of our model to qualitatively resemble the model of Lande (2014), in which species that invest more in plasticity fare better in extreme environments (e.g., contrast the blue and green species when $\varepsilon = 6$ in Fig. S1C). This assumption serves to increase the strength of selection on plasticity in variable environments.

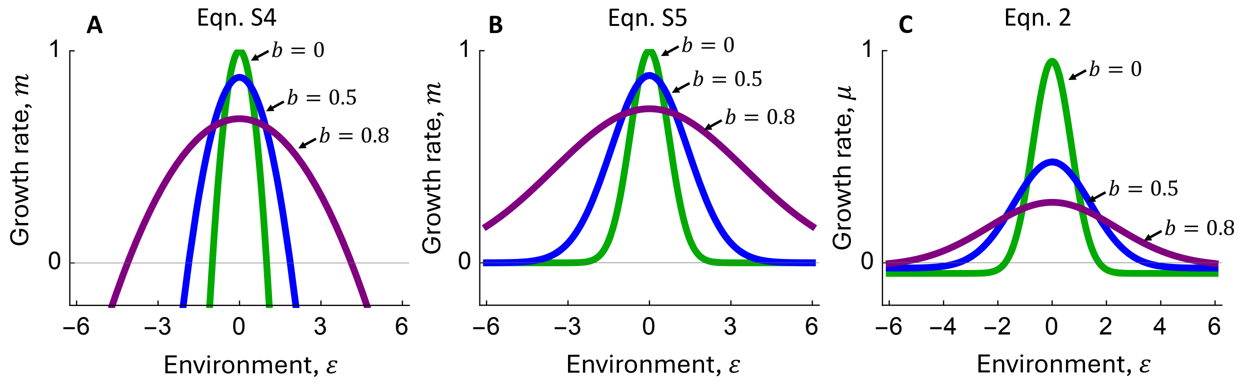


**Fig. S1 Environmental tolerance curves under different model structures. A** Effect of the environment on per-capita population growth rate for species that differ in investment in plasticity, $b$, under Eqn. S4. This takes $m_{max} = 1, \gamma = 2, \gamma_b = 1$, and $a = 0$. **B** As in panel **A**, but with growth rates determined by Eqn. S5. Parameter values here are $M_{max} = 1, \gamma = 2, \gamma_b = 1$, and $a = 0$. **C** Environmental tolerance curves based on Eqn. 2 of the main text, which was used in all analyses. Parameter values here are $\mu_{max} = 1, \gamma = 2, \gamma_b = 1$, and $a = 0$.

### S2 Supplementary Results

*S2.1 Effects of cost of plasticity on evolution of reaction norm slope*

In the absence of competitive interactions, increases in the cost of plasticity, $\gamma_b$, generally selected for species that invest less in phenotypic plasticity and, at a sufficiently high cost of plasticity $\gamma_b$, we found situations in which it was optimal to not invest in plasticity at all and instead specialize in high or low environmental values (Fig. S2). To understand this result, contrast the results for optimal reaction norm traits observed at the intermediate cost of plasticity $\gamma_b = 0.85$ – the value used in results presented in the main text (Fig. 2, Fig. S2D & E) – to results obtained with a lower cost of plasticity ($\gamma_b = 0.8$, Fig. S2A & D) and results in which there is a higher cost of plasticity ($\gamma_b = 0.9$, Fig. S2C & F). A relatively low cost of plasticity leads species to invest slightly more in plasticity at all amplitudes (contrast Fig. S2A to S2B). However, a relatively high cost of plasticity can lead to an abrupt change in the optimal reaction norm traits from a high investment in plasticity and no specialization, to a strategy that involves specializing in high or low environmental values (i.e., shifting $a_i$ so that growth is maximized during peaks or troughs in the environmental cycle; see amplitudes ~2-5 in Fig. S2C1-2). This occurs because forgoing investment in plasticity and instead specializing on peaks or troughs allows species to accrue large fitness benefits during those peaks or troughs (see orange or blue lines in Fig. S2F4) that ultimately outweighs the net fitness of the (now relatively costly) plastic strategy (see Fig. S2F1-2). The bimodality of the fitness landscape here (yielding two equally fit strategies) arises from the perfect symmetry of the environmental tolerance curves along with the bimodal distribution of the environment in sinusoidal cycles (Kremer & Klausmeier 2017).

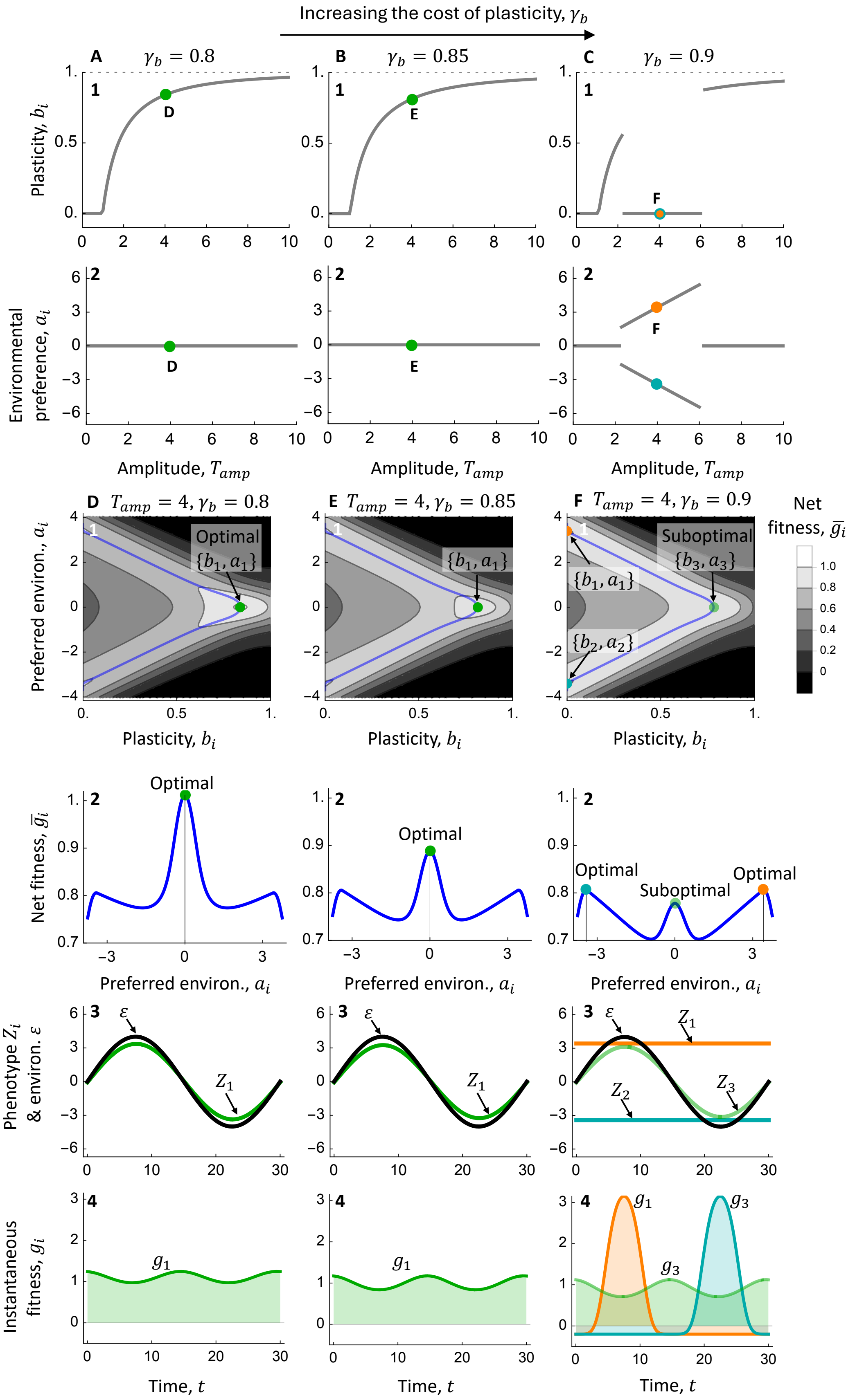

Increasing the cost of plasticity, $\gamma_b$
A $\gamma_b = 0.8$
B $\gamma_b = 0.85$
C $\gamma_b = 0.9$
Plasticity, $b_i$
Environmental preference, $a_i$
Amplitude, $T_{amp}$
D $T_{amp} = 4, \gamma_b = 0.8$
E $T_{amp} = 4, \gamma_b = 0.85$
F $T_{amp} = 4, \gamma_b = 0.9$
Optimal $\{b_1, a_1\}$
$\{b_1, a_1\}$
Suboptimal $\{b_3, a_3\}$
$\{b_2, a_2\}$
Preferred environ., $a_i$
Net fitness, $\bar{g}_i$
Optimal
Suboptimal
Phenotype $Z_i$ & environ. $\varepsilon$
$\varepsilon$
$Z_1$
$Z_2$
$Z_3$
Instantaneous fitness, $g_i$
$g_1$
$g_3$
Time, $t$

**Fig. S2 Increasing the cost of plasticity $\gamma_b$ generally reduces the optimal investment in phenotypic plasticity (set here by the reaction norm slope $b_i$).** Plasticity costs are varied from $\gamma_b = 0.8$ in **A & D** to $\gamma_b = 0.85$ in **B & E** to $\gamma_b = 0.9$ in **C & F. A-C** Panels 1 show the optimal reaction norm slope $b_i$, whereas panels 2 show the corresponding optimal preferred environment $a_i$. Here, the optimal reaction norm traits maximize the net fitness per period (when $N_i << K$). The dashed line indicates $b_i = 1$, corresponding to a reaction norm slope that matches changes in the optimal environment. The indicated points are considered in more detail in **D-F**. **D-F** Panels labeled 1 show the effects of preferred environment $a_i$ and reaction norm slope $b_i$ on the net fitness per period at an amplitude of 4. The colored points indicate the optimal traits, and the blue line traces a ridge across the fitness landscape (i.e., they show the values of $a_i$ that maximize fitness across a gradient in $b_i$). Panels labeled 2 show this trace of the fitness landscape (i.e., these panels give the net fitness per period across the values of $a_i$ that maximize fitness across a gradient in $b_i$). Panels labeled 3 show the coupled dynamics of the phenotype and environment for the optimal strategies, whereas panels labeled 4 show the associated fitness across the environmental cycle. Note that, in panels F3-4 the green species is suboptimal (i.e., its net growth rate per period is less than that of the orange and blue species) and is included here for comparison. Common parameter values are $\mu_{max} = 4, \gamma = 0.5, d_p = 0.05, \tau = 30$ and $\lambda = 10$.

We next consider how plasticity costs affect plasticity evolution in the presence of competition. When compared to findings based on an intermediate cost of plasticity ($\gamma_b = 0.85$, as in the main text Fig. 3, Fig. S3B & F-G), we found that a weakened cost of plasticity ($\gamma_b = 0.8$) sharply reduced the diversity of evolutionarily stable communities (Fig. S3A & D-E), whereas a relatively strong cost of plasticity ($\gamma_b = 0.9$) enhanced the diversity of evolutionarily stable communities at high amplitudes (Fig. S3C & H-I). When the cost of plasticity is relatively weak ($\gamma_b = 0.8$), evolutionarily stable communities comprise only a single species that invests relatively strongly in plasticity (Fig. S3A1-2; see also Fig. S2A-B). The relatively high investment in plasticity means that the plastic species fills sufficient niche space to competitively exclude alternative strategies – notice in Fig. S3D1-2 and Fig. S3E1-2 that specialists (i.e., species with $a_i$ larger or smaller than $0$) that do not invest in plasticity are unable to persist in the environment produced by the resident plastic species (Fig. S3D3-4 and Fig. S3E3-4). The opposite result occurs when the cost of plasticity is relatively strong ($\gamma_b = 0.9$). A strong cost of plasticity causes the plastic species to invest relatively weakly in plasticity (Fig. S3C1; see also Fig. S2B-C), which means that it does not fill as much niche space (Fig. S3H3-4 and Fig. S3H3-4) and, as such, the two specialist species that do not invest in plasticity (the orange and blue species in Fig. S3C, H-I) are able to coexist with the plastic species over a wider range of amplitudes (contrast Fig. S3B with Fig. S3C). These results show that the cost of plasticity modulates community diversity by controlling the amount of niche space captured by the plastic species in evolutionarily stable communities.

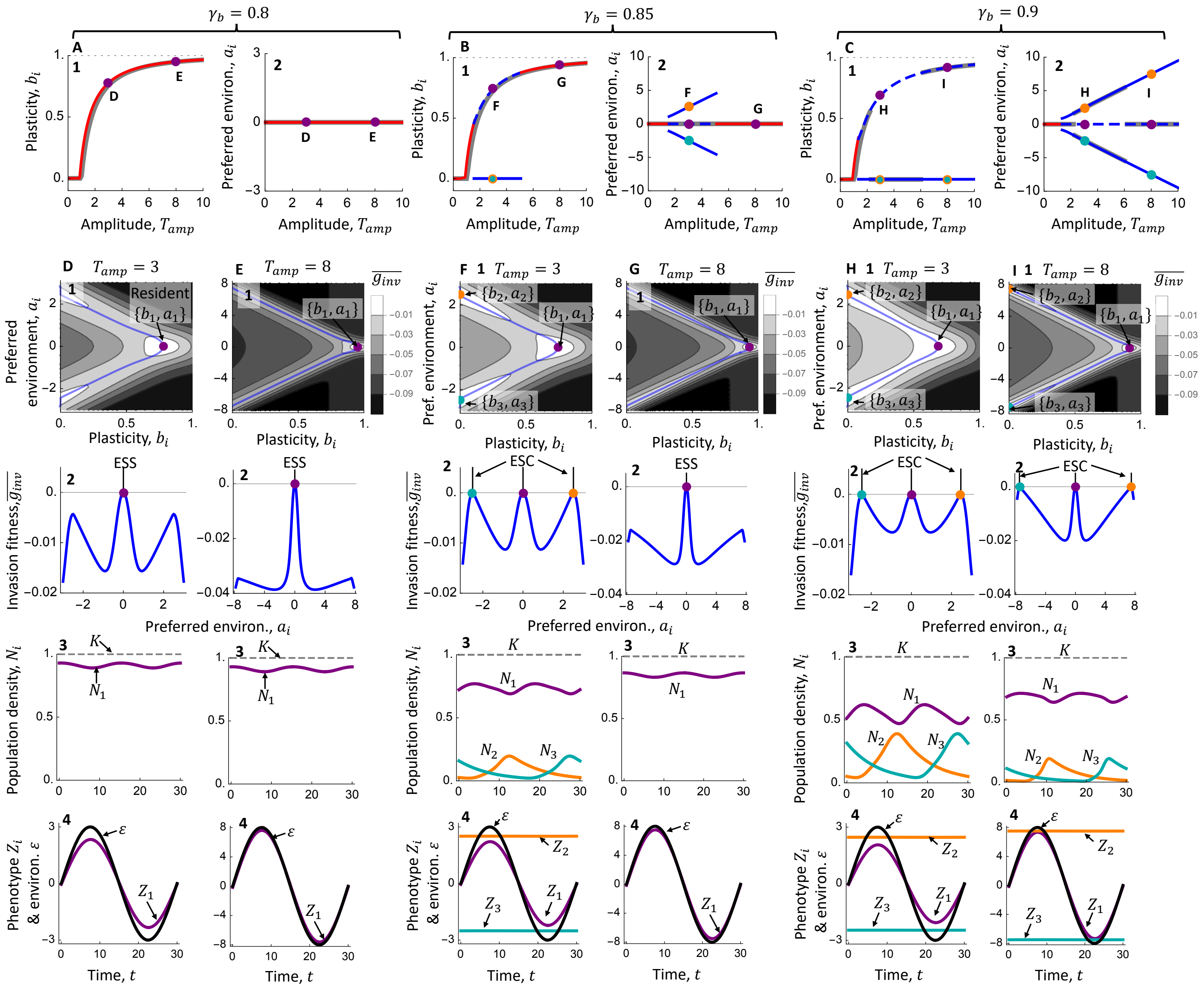
$\gamma_b = 0.8$
$\gamma_b = 0.85$
$\gamma_b = 0.9$
Plasticity, $b_i$
Preferred environ., $a_i$
Amplitude, $T_{amp}$
$T_{amp} = 3$
$T_{amp} = 8$
Resident
ESS
ESC
Invasion fitness, $\overline{g_{inv}}$
Population density, $N_i$
Phenotype $Z_i$ & environ. $\varepsilon$
Time, $t$

189

**Fig. S3 Effects of increasing the cost of plasticity $\gamma_b$ on the response of evolutionarily stable reaction norm traits to amplitude (varying $\gamma_b$ from $0.8$ in A & D-E, to $0.85$ in B & F-G, to $0.9$ in C & H-I). A-C** Evolutionarily stable communities (ESCs) comprising species with sets of reaction norm traits $\overrightarrow{\{a, b\}}$ across amplitudes of environmental fluctuation. Results based on $\gamma_b = 0.85$ are equivalent to those in Figs. 2-3 of the main text and are included here to aid comparison. Panels 1 show the evolutionarily stable investment in plasticity $b_i$, whereas panels 2 show the corresponding environmental preferences $a_i$. The color of ESCs are varied with the species richness of the community, and the line types change in multispecies ESCs to map the traits in 1 to those in 2. The gray line shows the optimal traits found in the absence of competition (Fig. S2, notice that the ESCs tend to be on top of the optimal strategies). The points indicated in these panels are examined in more detail in **D-I**. **D-I** These panels show details about ESCs found at an amplitude of 3 or 8 for the respective cost of plasticity ($\gamma_b = 0.8$ D-E, $\gamma_b = 0.85$ in F-G, and $\gamma_b = 0.9$ in H-I). Panels labeled 1 show the invasion fitness for rare mutants with traits $b_i$ and $a_i$ at the ecological attractor set by the resident species or community with traits indicated by the colored points. As in Fig. S2D1-F1, the blue line traces a ridge across the fitness landscape (i.e., they show the values of $a_i$ that maximize invasion fitness across a gradient in $b_i$). Panels labeled 2 show this trace of the fitness landscape (i.e., these panels give the net invasion fitness along the values of $a_i$ that maximize fitness across values of $b_i$). Note that the resident species have a net fitness of zero because they are at their ecological attractor (see Eqn. 5). Panels labeled 3 show the ecological attractors for the ESCs, and panels labeled 4 show the phenotype-environment dynamics for the ESCs. Common parameter values are $\mu_{max} = 4, \gamma = 0.5, d_p = 0.05, d = 0.1,\ K = 1, \tau = 30$ and $\lambda = 10$.

*S2.2 The optimal acclimation rate in the absence of competition.*

The optimal investment in plasticity $\lambda_i$ exhibited a unimodal relationship to the amplitude of environment fluctuations (Fig. S4A). At low amplitudes, increasing environmental fluctuation amplitude $T_{amp}$ – e.g., from $T_{amp} = 1.5$ (Fig. S4C) to $T_{amp} = 4$ (Fig. S4D) – favored species that invest more in plasticity $\lambda_i$ in order to avoid the relatively large phenotype-environment mismatches at higher amplitudes. For example, notice that the species with the optimal traits at $T_{amp} = 1.5$ (i.e., the green species in Fig. S4C and S4D3) experiences relatively large phenotype-environment mismatches at $T_{amp} = 4$ and is therefore less fit than species that invest more in plasticity (e.g., the purple species in Fig. S4D). At higher amplitudes (e.g., $T_{amp} > {\sim}6$), species that invested less in plasticity and instead shifted their environmental preferences $a_i$ were favored (Fig. S4A-B, E). These species (e.g., the orange and blue species in Fig. S4E) accumulate relatively large fitness benefits during periods of time yielding phenotype-environment matches (e.g., times 0-10 or 10-20, respectively, for the orange and blue species in Fig. S4E3).

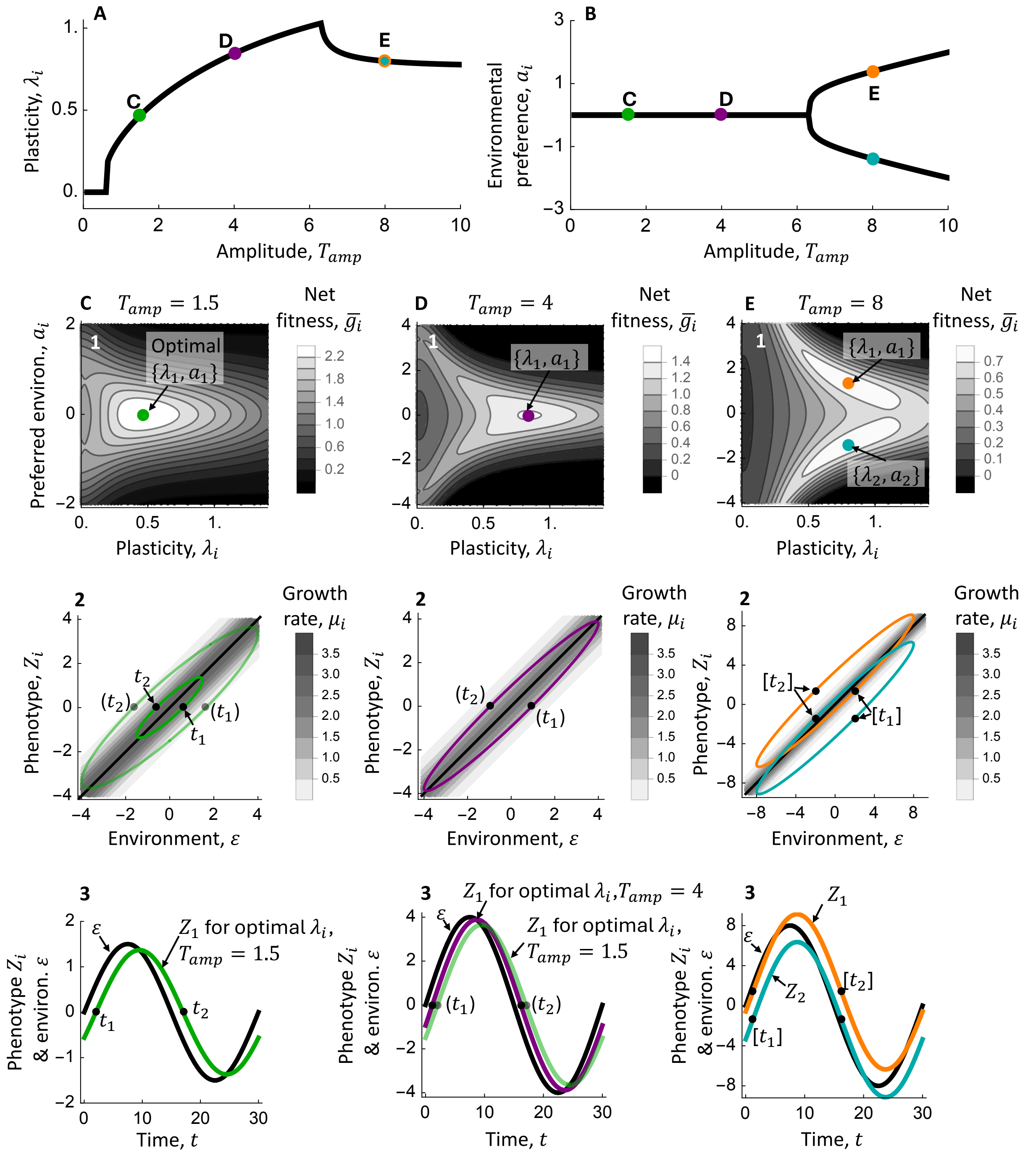


**Fig. S4 Effects of the amplitude of environmental fluctuations on the optimal traits when the acclimation rate controls investment in plasticity and competition is absent. A-B** The optimal acclimation rate $\lambda_i$ along with the associated optimal preferred environment $a_i$ (these are identical to gray lines shown in Fig. 4A-B of the main text). The optimal traits maximize the net fitness per period (when $N_i << K$). The indicated points are considered in more detail in **C-E**. **C-E** Panels labeled 1 show the effects of investment in plasticity and the preferred environment $a_i$ on the net fitness per period at the indicated amplitude $T_{amp}$. The colored points show the optimal traits. Panels labeled 2 show the phenotype dynamics (colored lines) overlain on the growth rate $\mu_i$ contour plots. In **C**, we also show the phenotype dynamics for this species when it is at $T_{amp} = 4$ (with the slightly transparent green line), to show that it experiences large phenotype-environment mismatches at that $T_{amp}$. Note that the cost of plasticity leads the maximum growth rate $\mu_i$ to be lower for species that invest more in plasticity (i.e., $\mu_i$ is larger in **E** than **D**, and $\mu_i$ in **D** is in turn larger than that in **C**). The points $t_{1,2}$, $(t_{1,2})$, and $[t_{1,2}]$ indicate sets of phenotypes and environments at two time points during cycles of amplitude $T_{amp} = 1.5$, $T_{amp} = 4$, and $T_{amp} = 8$, respectively, shown in panels labeled 3. Panels labeled 3 show phenotype-environment dynamics for the optimal species at the respective amplitude. The phenotype dynamics for the optimal species at $T_{amp} = 1.5$ (green) are also displayed at $T_{amp} = 4$ in **D**, to demonstrate the relatively large phenotype-environment mismatches that come from reduced investment in plasticity at this amplitude. Parameter values are $\mu_{max} = 4$, $\gamma = 0.5$, $\gamma_b = 0.85$, $d_p = 0.05$, $\tau = 30$ and $b = 1$.

*S2.3 Effects of cost of plasticity on evolution of acclimation rate*

Increasing the cost of plasticity (set by the acclimation rate $\lambda_i$) does not qualitatively alter evolutionarily stable traits across amplitudes, but it does reduce investment in plasticity and shift the occurrence of major bifurcation events (i.e., gain or loss of species) to lower amplitudes (Fig. S5). To understand this, contrast findings based on an intermediate cost of plasticity ($\gamma_b = 0.5$, Fig. S5B & F-G; note that this is identical to results shown in the main text Fig. 4) to those observed with a weak cost of plasticity ($\gamma_b = 0.3$, Fig. S5A & D-E) and with a relatively strong cost of plasticity ($\gamma_b = 0.7$, Fig. S5C & H-I). A relatively weak cost of plasticity favored species with increased investment in phenotypic plasticity (Fig. S5A), as species do not need to sacrifice as much fitness to reap the rewards of investing in plasticity. This means that the plastic species is better able to competitive exclude alternative strategies by more efficiently capturing the available niche space, which ultimately leads to increases in the amplitude at which the system diversifies (i.e., from an $T_{amp}$~4 when $\gamma_b = 0.5$ to $T_{amp}$~8 when $\gamma_b = 0.3$). The opposite patterns occur at relatively high costs of plasticity ($\gamma_b = 0.7$), where evolution favored species with reduced investment in phenotypic plasticity (notice that $\lambda_i$ is less than 1 across high amplitudes in Fig. S5C), as the rewards of investing in plasticity are offset by increased fitness costs. This reduction in investment in plasticity reduced the capacity for one plastic species to competitively exclude alternative strategies, and thereby led to reductions in the amplitude at which the community diversifies (i.e., to $T_{amp}$~3 when $\gamma_b = 0.7$).

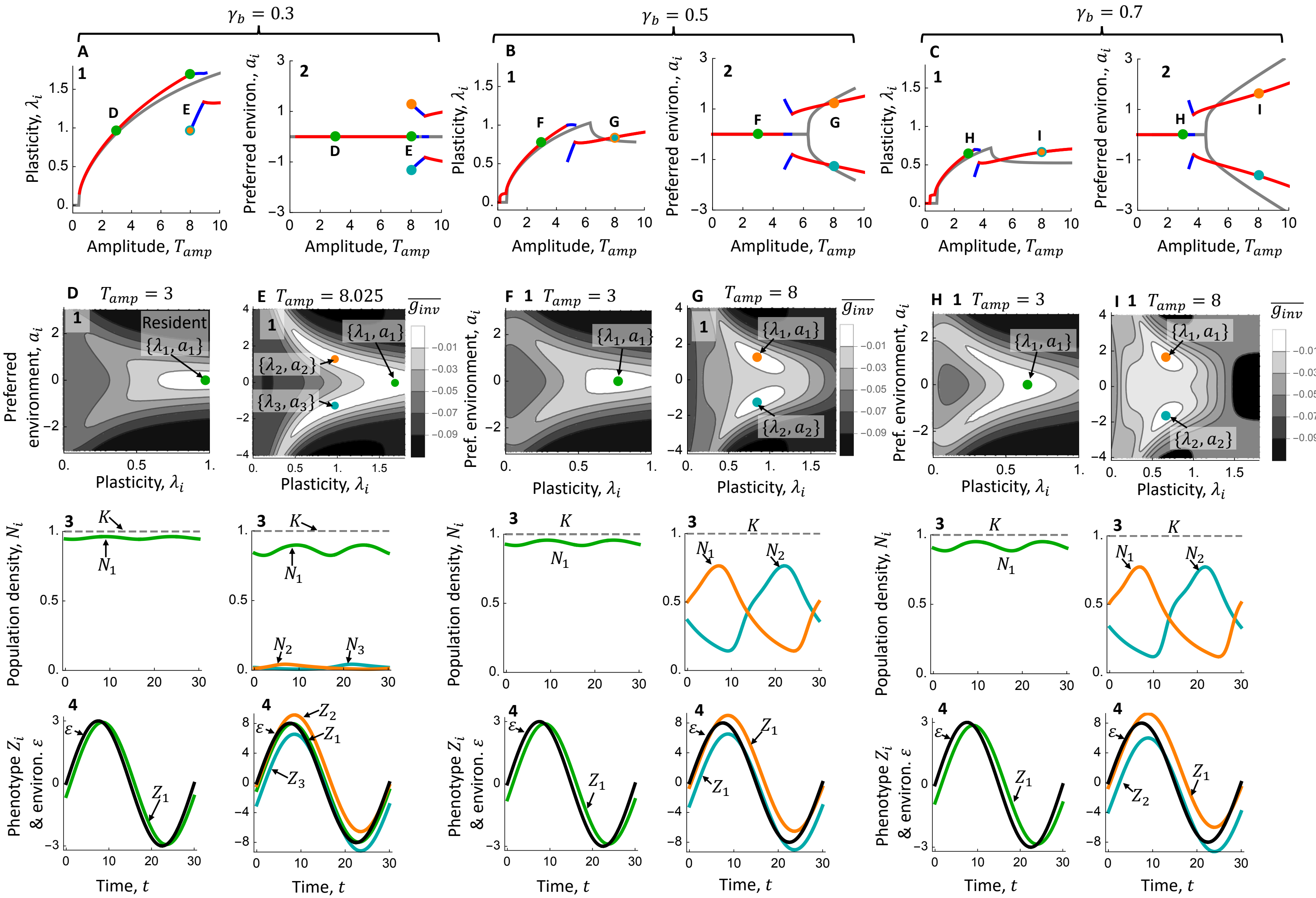

$\gamma_b = 0.3$
$\gamma_b = 0.5$
$\gamma_b = 0.7$
A
B
C
1
2
Plasticity, $\lambda_i$
Preferred environ., $a_i$
Amplitude, $T_{amp}$
D
E
F
G
H
I
D 1 $T_{amp} = 3$
E 1 $T_{amp} = 8.025$
F 1 $T_{amp} = 3$
G 1 $T_{amp} = 8$
H 1 $T_{amp} = 3$
I 1 $T_{amp} = 8$
Resident
$\{\lambda_1, a_1\}$
$\{\lambda_2, a_2\}$
$\{\lambda_3, a_3\}$
$\overline{g_{inv}}$
−0.01
−0.03
−0.05
−0.07
−0.09
Preferred environment, $a_i$
Pref. environment, $a_i$
3
$K$
$N_1$
$N_2$
$N_3$
Population density, $N_i$
4
$\varepsilon$
$Z_1$
$Z_2$
$Z_3$
Phenotype $Z_i$ & environ. $\varepsilon$
Time, $t$

**Fig. S5 Increasing the cost of plasticity $\gamma_b$ affects the response of evolutionarily stable communities to amplitude (varying $\gamma_b$ from $0.3$ in A & D-E, to $0.5$ in B & F-G, to $0.7$ in C & H-I). A-C** ESCs consisting of species with traits $\overrightarrow{\{a, \lambda\}}$ across amplitudes of environmental fluctuation. Results based on $\gamma_b = 0.5$ are identical to the findings in Fig. 4 of the main text and are included here for reference. Panels 1 show the evolutionarily stable investment in plasticity rate $\lambda_i$, whereas panels 2 show the species' associated environmental preferences $a_i$. We vary the color of ESCs with the species richness of the community. For the 3 species ESCs, the species that invests more in plasticity always has $a_i = 0$. The gray line shows the optimal traits found in the absence of competition. The points indicated in these panels are the subject of detailed examination in **D-I**. **D-I** These panels display details regarding ESCs found at amplitude of 3 or 8 for the respective cost of plasticity ($\gamma_b = 0.3$ D-E, $\gamma_b = 0.5$ in F-G, and $\gamma_b = 0.7$ in H-I). All panels labeled 1 give the net invasion fitness for rare mutants with traits $\lambda_i$ and $a_i$ in the attractor of the resident species or community. These residents' trait values are indicated by the colored points. Note that the resident species have a net fitness of zero because they are at their ecological attractor (see Eqn. 5), and all other trait values have a negative invasion fitness (and hence these are ESCs). Panels labeled 2 show the ecological attractors for the or ESCs, and panels labeled 3 show the dynamics of phenotypes and the environment for the ESCs. Common parameter values are $\mu_{max} = 4, \gamma = 0.5, d_p = 0.05, d = 0.1,\ K = 1, \tau = 30$ and $b = 1$.

*S2.4 Effects of the cost of plasticity on community properties*

Accompanying our investigations above of the effects of cost of plasticity on optimal trait values for plastic capacity, $b$, and rate, $\lambda$, in the presence of competition, we also illustrate the corresponding consequences for community-level properties (as described in the main text) for $b$ (Fig S6) and $\lambda$ (Fig. S7). In general, as plasticity becomes more costly, communities show lower, more temporally variable total biomass.

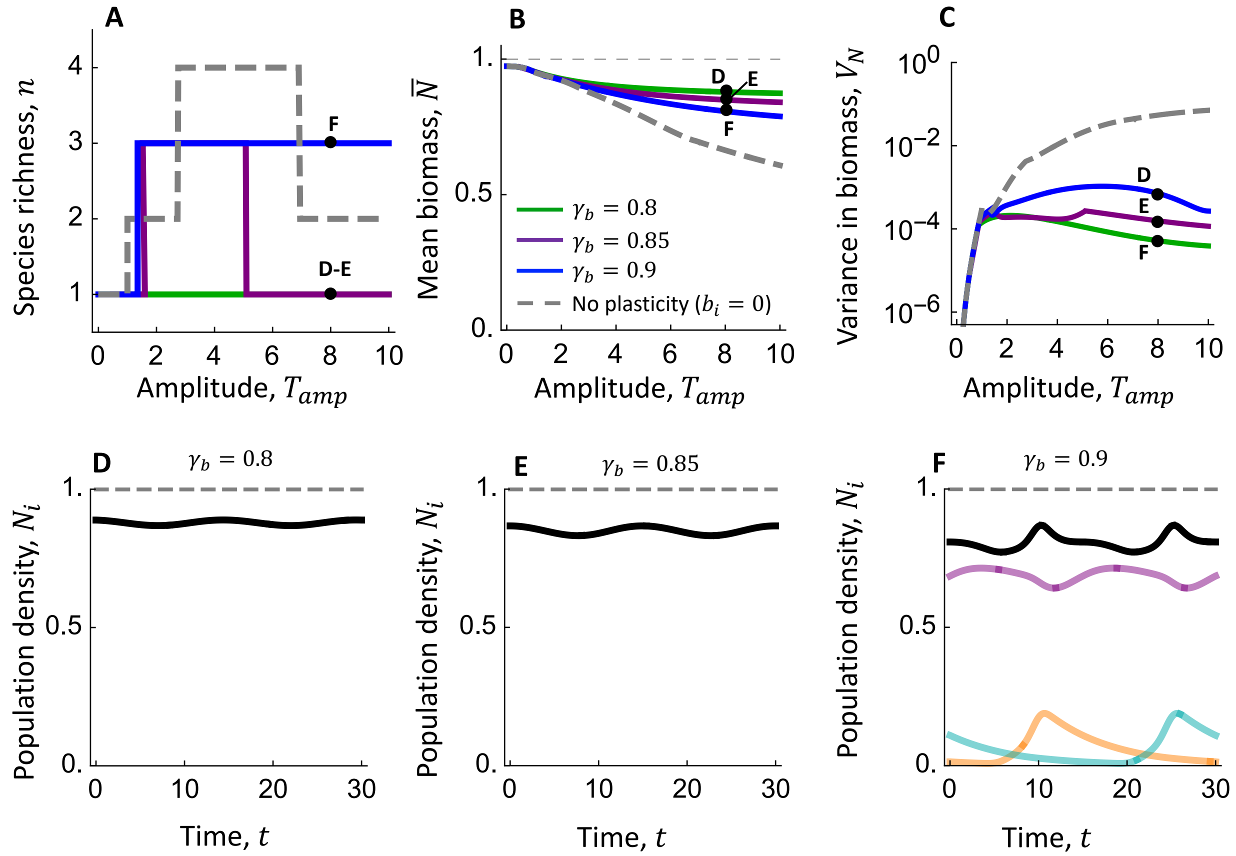


**Fig. S6 Increasing the cost of plasticity – where here plasticity is controlled by the reaction norm slope $b_i$ – increases species richness (A), reduces the mean community density (B, Eqn. 7), and increases the temporal variance in community density (C, Eqn. 8). A-C** These panels show the respective community property for the ESCs obtained for three values of the costs of plasticity: $\gamma_b = 0.8$ (green line), $\gamma_b = 0.85$ (purple line; note that these results are identical to those in Fig. 6 of main text), and $\gamma_b = 0.9$ (blue line). For reference, we also include results obtained in the absence of plasticity (gray, dashed line, Fig. 5A). The black points show communities that are the subject of more detailed examination in **D-F. D-F** Population and community dynamics at $T_{amp} = 8$ for the three values of the costs of plasticity. In each panel, the black line is the total density of the community, the semi-transparent colored lines show population dynamics for individual species (note that there is only one species in **D-E**), and the gray dashed line is the carrying capacity.

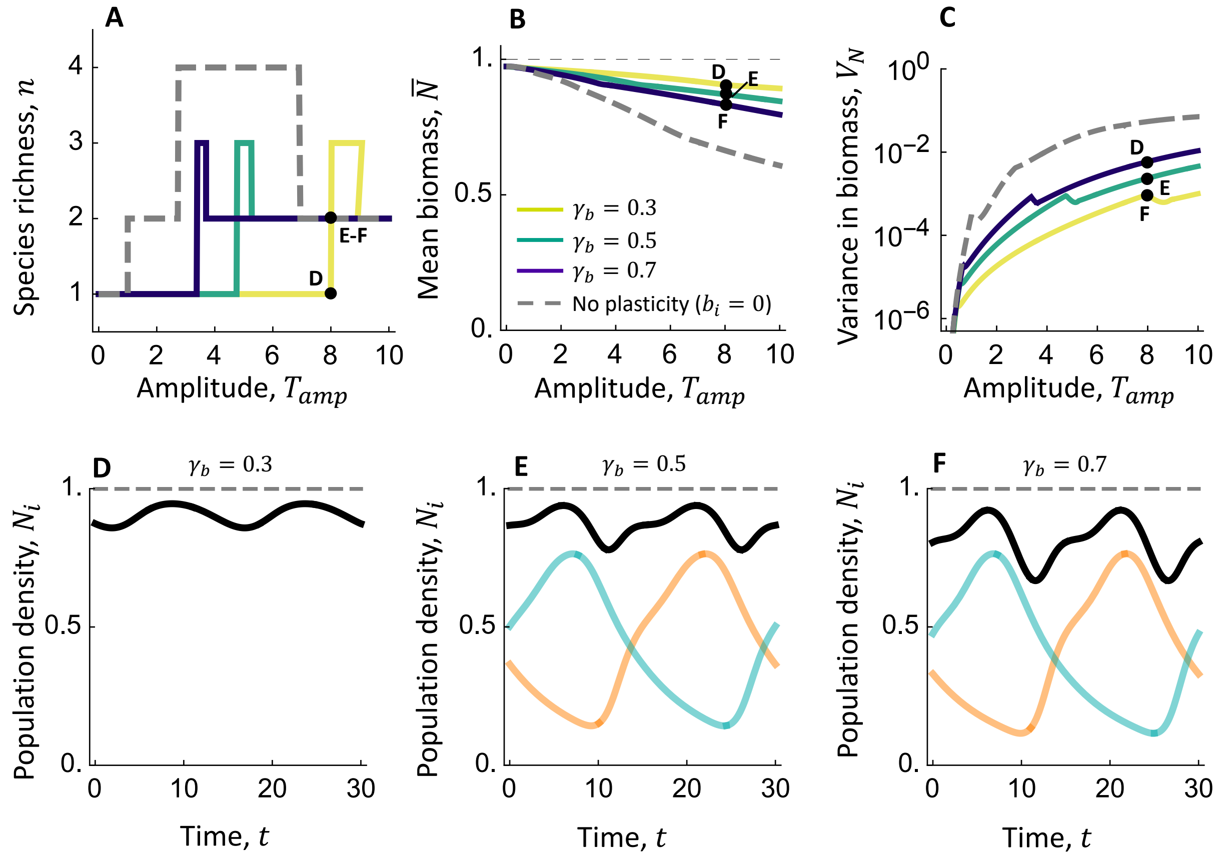


**Fig. S7 When plasticity is determined by the acclimation rate $\lambda_i$, increases in the cost of plasticity $\gamma_b$ alters the patterning of species richness across amplitudes (A), reduces the mean community density (B, Eqn. 7), and increases the temporal variance in community density (C, Eqn. 8). A-C** These panels show the respective community property for the ESCs obtained for three values of the costs of plasticity: $\gamma_b = 0.8$ (green line), $\gamma_b = 0.85$ (purple line; note that these results are identical to those in Fig. 6 of main text), and $\gamma_b = 0.9$ (blue line). For reference, we also include results obtained in the absence of plasticity (gray, dashed line, Fig. 5A). The black points show communities that are the subject of more detailed examination in **D-F. D-F** Population and community dynamics at $T_{amp} = 8$ for the three values of the costs of plasticity. In each panel, the black line is the total density of the community, the semi-transparent colored lines show population dynamics for individual species (note that there is only one species in **D-E**), and the gray dashed line is the carrying capacity.
